# The Nucleotide Transformer: Building and Evaluating Robust Foundation Models for Human Genomics

**DOI:** 10.1101/2023.01.11.523679

**Authors:** Hugo Dalla-Torre, Liam Gonzalez, Javier Mendoza-Revilla, Nicolas Lopez Carranza, Adam Henryk Grzywaczewski, Francesco Oteri, Christian Dallago, Evan Trop, Bernardo P. de Almeida, Hassan Sirelkhatim, Guillaume Richard, Marcin Skwark, Karim Beguir, Marie Lopez, Thomas Pierrot

**Affiliations:** InstaDeep; Nvidia; TUM

**Author notes:** Equal supervision (Marie Lopez, Thomas Pierrot). Corresponding author(s) &.

## Abstract

Closing the gap between measurable genetic information and observable traits is a longstanding challenge in genomics. Yet, the prediction of molecular phenotypes from DNA sequences alone remains limited and inaccurate, often driven by the scarcity of annotated data and the inability to transfer learning between prediction tasks. Here, we present an extensive study of foundation models pre-trained on DNA sequences, named the Nucleotide Transformer, ranging from 50M up to 2.5B parameters and integrating information from 3,202 diverse human genomes, as well as 850 genomes selected across diverse phyla, including both model and non-model organisms. These transformer models yield transferable, context-specific representations of nucleotide sequences, which allow for accurate molecular phenotype prediction even in low-data settings. We show that the developed models can be fine-tuned at low cost and despite low available data regime to solve a variety of genomics applications. Despite no supervision, the transformer models learned to focus attention on key genomic elements, including those that regulate gene expression, such as enhancers. Lastly, we demonstrate that utilizing model representations can improve the prioritization of functional genetic variants. The training and application of foundational models in genomics explored in this study provide a widely applicable stepping stone to bridge the gap of accurate molecular phenotype prediction from DNA sequence. Code and weights available on GitHub in Jax and HuggingFace in Pytorch. Example notebooks to apply these models to any downstream task are available on HuggingFace.

## Introduction

Foundation models in artificial intelligence (AI) are characterized by their large-scale nature, incorporating millions of parameters trained on extensive datasets. These models can be adapted for a wide range of subsequent predictive tasks and have profoundly transformed the AI field. Notable examples in natural language processing (NLP) include the so-called language models (LMs) BERT [1] and GPT [2]. LMs have gained significant popularity in recent years owing to their ability to be trained on unlabeled data, creating general-purpose representations capable of solving downstream tasks. One way they achieve a comprehensive understanding of language is by solving billions of cloze tests, in which they predict the correct word to fill in the blank in a given sentence. This approach is known as masked language modeling [1]. Early instances of foundation models applying this objective to biology involved training LMs on protein sequences, where they were tasked with predicting masked amino acids in large protein sequence datasets [3–5]. These protein LMs, when applied to downstream tasks using transfer learning, demonstrated the ability to compete with and even outperform previous methods for tasks such as predicting protein structure [3, 4] and function [6, 7], even in data scarce regiments [8].

Beyond protein sequences, the dependency patterns encoded in DNA sequences play a fundamental role in understanding genomic processes, from characterizing regulatory regions to assessing the impact of individual variants within their haplotypic context. In this context, specialized deep learning (DL) models have been trained to uncover meaningful patterns of DNA [9–12]. For example, DL models have been used to predict gene expression from DNA sequences [13–18], with recent advancements combining convolutional neural networks (CNN) and transformer architectures enabling the encoding of regulatory elements located up to 100 kilobases (kb) upstream [19]. The abundance of data generated by modern genomics research presents both an opportunity and a challenge. On one hand, intricate patterns of natural variability across species and populations are readily available; on the other hand, powerful deep learning methods capable of handling large-scale data are necessary for accurate signal extraction from unlabeled datasets. Large foundation models trained on nucleotides sequences appear to be a natural choice for addressing this challenge [20–26].

With the Nucleotide Transformer we present a systematic study and benchmark on how to construct and evaluate robust foundation models to encode genomic sequences. We initiated our study by building four distinct language models of varying sizes, ranging from 500M up to 2.5B parameters. These models were pre-trained on three different datasets including the human reference genome, a collection of 3,202 diverse human genomes, and 850 genomes from various species. After training, we leveraged the representations (i.e. embeddings) from each of these models in two fashion. To evaluate the Nucleotide Transformer’s stability in performance while adapting to various tasks, we trained each model on a diverse set of 18 genomic curated prediction tasks and compared them with three alternative DNA foundational models, alongside one state-of-the-art non-foundational model, using a systematic 10-fold cross-validation procedure. Moreover, to broaden our evaluation, we compared our best-performing model and three state-of-the-art supervised baseline models that have been optimized for the specific tasks at hand. To decipher the sequence features learned during pre-training, we explored the models’ attention maps, perplexities, and conducted data dimensionality reduction on their embeddings. Additionally, we evaluated the embeddings’ capacity to model the impact of functionally important genetic variants in humans, through zero-shot-based scores. Expanding upon the findings from the initial set of experiments, we developed a second set of four language models, with decreasing sizes from 500M to 50M parameters, to investigate the scaling laws of such models. We successfully constructed a model that achieves the performance of our previous best model with only one-tenth the number of parameters and while doubling the perception field size.

## Results

### The Nucleotide Transformer model accurately predicts diverse genomics tasks

We developed a collection of transformer-based DNA language models, dubbed Nucleotide Transformer (NT), which have learned general nucleotide sequence representations from 6kb unannotated genomic data (Fig. 1a; Methods). Inspired by trends in NLP, where larger training datasets and model sizes have demonstrated improved performance [27], we constructed transformer models with varying parameter sizes and datasets: (i) a 500 million parameter model trained on sequences extracted from the human reference genome (Human ref 500M), (ii) a 500 million and (iii) a 2.5 billion parameter model both trained on 3,202 genetically diverse human genomes[28] (1000G 500M and 1000G 2.5B respectively), and (iv) a 2.5 billion parameter model, encompassing 850 species from diverse phyla (Multispecies 2.5B), including 11 model organisms (Fig. 1c; Supplementary Tables 1, 2, 3, 4).

**Figure 1.**
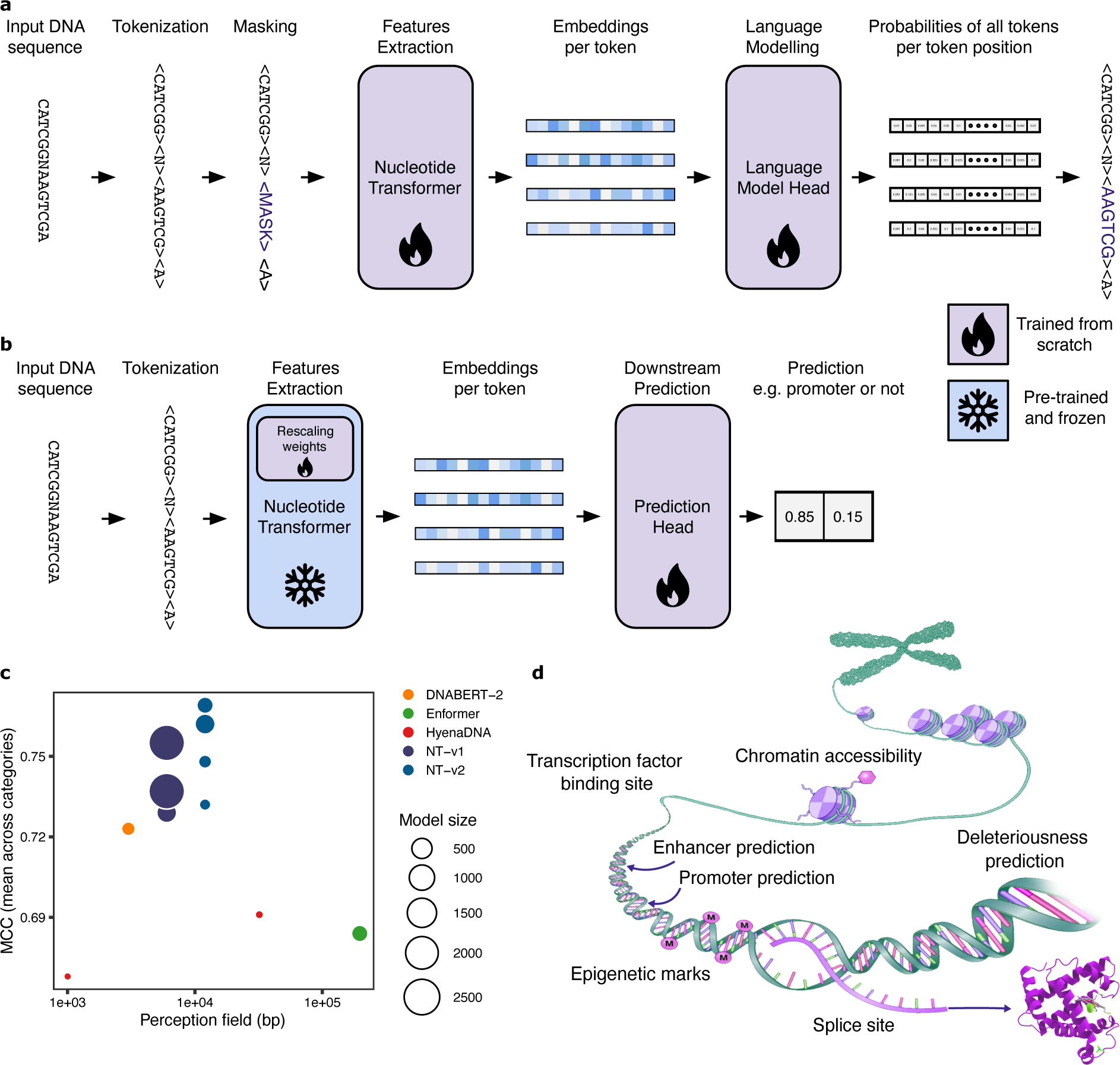
The Nucleotide Transformer: effective methodology to pre-train, fine-tune, analyse and compare foundational models for genomics. **a,b)** Overview of the Nucleotide Transformer training **(a)** and application for downstream genomic prediction tasks through fine-tuning **(b)**. Downstream task prediction through probing is similar but without the rescaling weights in the Nucleotide Transformer. **c)** Comparison of the Nucleotide Transformer models to other foundational genomics models in terms of perception field size, number of parameters and performance over our benchmark made of 18 curated downstream tasks. **d)** Graphical representation of genomic features considered for downstream tasks.

In order to evaluate the efficacy of these models in predicting varied molecular phenotypes, we curated 18 genomic datasets from publicly available resources encompassing splice site prediction tasks, (GENCODE [29]), promoter tasks (Eukaryotic Promoter Database [30]), and histone modification and enhancer tasks (ENCODE [31–33]), each designed to be of reasonable size to enable fast and rigorous cross-validation procedures (Fig. 1d; Supplementary Table 5, see Methods). While larger datasets for supervised models are available, this compilation of 18 genomic datasets offers a diverse and robust selection for scrutinizing the models’ adaptability across numerous tasks in a statistically rigorous manner and for comparison with other DNA self-supervised foundational models. These datasets were processed into a standardized format to facilitate experimentation and ensure reproducibility in the assessment of large language model performance (see Methods). We evaluated the transformer models through two different techniques: probing and fine-tuning (Fig. 1b). Probing refers to the use of learned LM embeddings of DNA sequences as input features to simpler models for predicting genomic labels. Specifically, we probed ten arbitrarily chosen layers of the LMs using either a logistic regression or a small multi-layer perceptron (MLP) composed of up to two hidden layers. In the case of fine-tuning, the LM head is substituted with either a classification or regression head, and a parameter-efficient technique is used for retraining (Methods). To ensure a fair and accurate comparison between different models, we implemented a 10-fold cross-validation strategy.

To compare our pre-trained foundational model schemes with standard, supervised methods in the field, we trained different variants of the BPNet convolutional architecture [9] from scratch on each of the 18 tasks (see Methods). The BPNet architecture has been used widely in genomics and represents a very strong default architecture for modelling small-sized datasets from scratch through supervised learning. We observed strong performance across tasks for the original BPNet model (average Matthews correlation coefficient (MCC) of 0.665), that we could improve by increasing its size to 28 million parameters (average MCC of 0.683), confirming that directly supervised convolutional architectures perform very well for genomics tasks (Fig. 2a,b). We next evaluated how probing and fine-tuning of the NT models compare with these supervised baseline models on our benchmark datasets. We considered the models to be equivalent to or better than other models if the resulting two standard deviations either overlapped or were superior to the reported baseline value, respectively.

**Figure 2.**
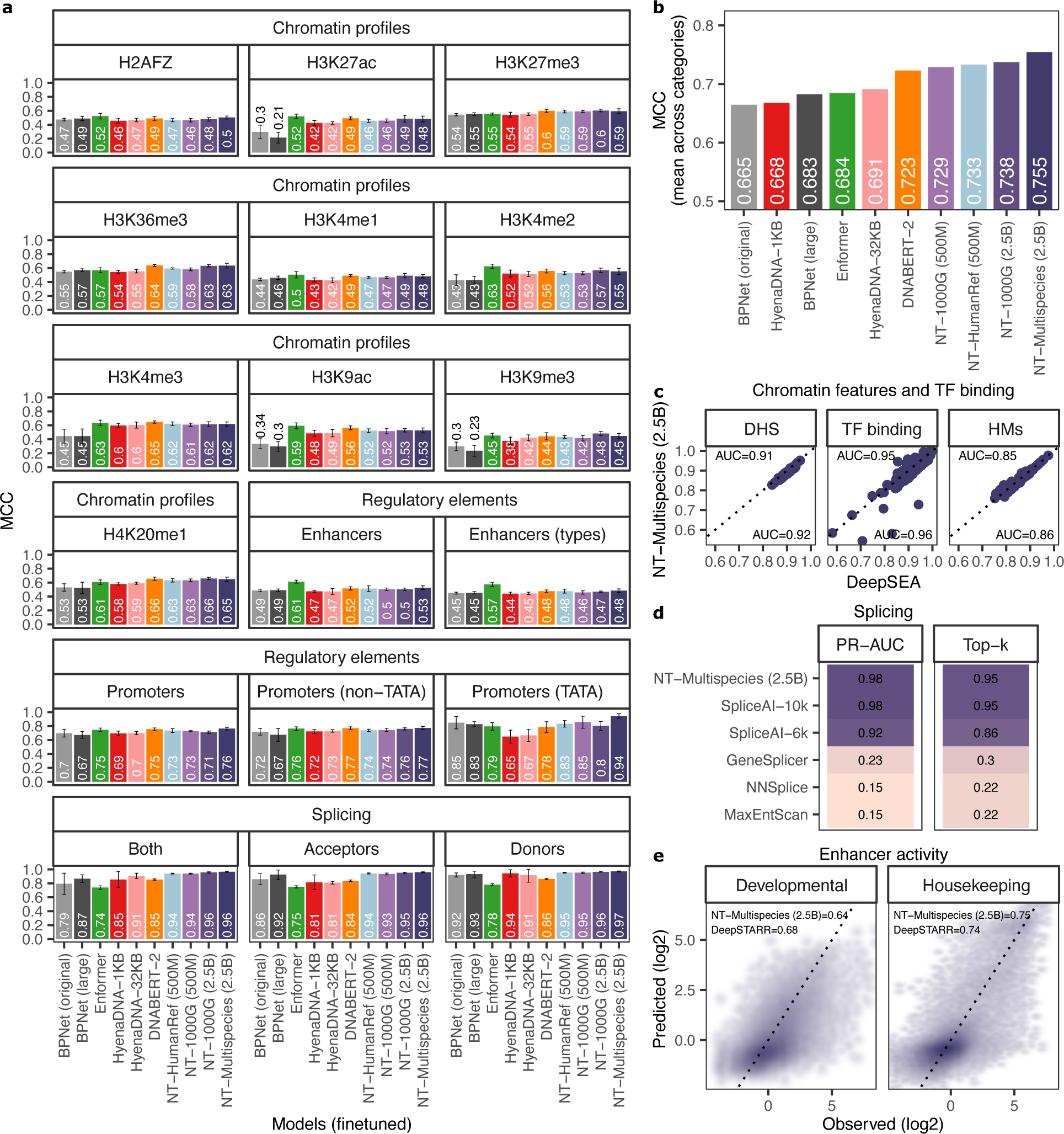
The Nucleotide Transformer model accurately predicts diverse genomics tasks after fine-tuning. **a)** Performance results across downstream tasks for fine-tuned NT models as well as HyenaDNA, DNABERT and Enformer pre-trained models based on the Matthew’s correlation coefficient (MCC). We also trained BPNet models from scratch for comparison (original: 121k parameters; large: 28M parameters). Error bars represent 2 SDs derived from 10-fold cross-validation. **b)** Normalized mean of MCC performance across downstream tasks (divided by category) for all language models after fine-tuning. **c)** The Multispecies 2.5B model performance on DNase I hypersensitive sites (DHS), histone marks (HMs), and transcription factor sites predictions from different human cells and tissues compared with the baseline DeepSEA model. Each dot represents the area under the ROC curve (AUC) for a different genomic profile. The average AUC per model is labeled. **d)** The Multispecies 2.5B model performance on predicting splice sites from the human genome, compared with the SpliceAI and other splicing models. **e)** The Multispecies 2.5B model performance on developmental and housekeeping enhancer activity predictions from *Drosophila melanogaster* S2 cells, compared with the baseline DeepSTARR model.

Using this criterion, NT models matched the baseline BPNet models in 5 tasks and exceeded them in 8 out of the 18 tasks through probing alone (Supplementary Fig.1 and Table 6), and significantly outperformed probing from raw tokens. In agreement with recent work [34], we observed that the best performance is both model- and layer-dependent (Supplementary Table 8). We also noted that the highest model performance is never achieved by using embeddings from the final layer, as shown in earlier work [5]. For instance, in the enhancer types prediction task, we observed a relative difference as high as 38% between the highest and lowest performing layer, indicating significant variation in learned representations across the layers (Supplementary Fig. 3). Compared to our probing strategy, our fine-tuned models either matched (6) or surpassed (12) out of the 18 baseline models (Fig. 2a,b and Table 7). Notably, finetuned NT models outperformed the probed models, and the larger and more diverse models consistently outperformed their smaller counterparts (Fig. 2a,b and Supplementary Table 9 and 7). These results support the necessity for fine-tuning the NT foundation models towards specific tasks to achieve superior performance. Our results also suggest that training on a diverse dataset, represented by the Multispecies 2.5B model, outperforms or matches the 1000G 2.5B model on several tasks derived from human-based assays (Fig. 2a,b). This implies that a strategy of increased diversity, rather than just increased model size, may lead to improved prediction performance, particularly when computational resources are limited.

Fine-tuning has not been extensively explored in previous work [5], possibly due to its demanding computational requirements. We overcame this limitation by adopting a recent parameter-efficient fine-tuning technique [35] which requires only 0.1% of the total model parameters (Fig. 1b; Methods). This approach allowed for faster fine-tuning on a single GPU, reduced storage needs by 1,000-fold over all fine-tuning parameters, while still delivering comparable performance. In practice, we observed that rigorous probing was slower and more computationally intensive than fine-tuning, despite the apparent simplicity of using straightforward downstream models on embeddings. This discrepancy arises from the significant impact of factors such as layer choice, downstream model selection, and hyperparameters on performance. Additionally, fine-tuning exhibited a smaller variance in performance, enhancing the robustness of the approach. Overall, this general approach is versatile and adaptable to various tasks without requiring adjustments to the model architecture or hyperparameters. This stands in contrast to supervised models, which typically feature distinct architectures and necessitate training from scratch for each task.

Finally, we aimed to evaluate the potential for large language DNA models to compete with robust baselines trained in a supervised manner using extensive datasets and optimized architectures. To do so, we applied the Multispecies 2.5B model to three additional genomic prediction tasks, which encompassed the classification of 919 chromatin profiles from a diverse set of human cells and tissues [10], predicting canonical splice acceptor and donor sites across the whole human genome [36], and predicting developmental and housekeeping enhancer activities from *Drosophila melanogaster* S2 cells [12] (Methods). Remarkably, the Multispecies 2.5B model achieved performance levels closely aligned with those of specialized deep learning models, despite no additional changes or optimizations of its original fine-tuning architecture. For instance, in the case of classifying chromatin feature profiles, we obtained area under the curve (AUC) values that were, on average, only approximately ~1% lower than those achieved by DeepSEA (Fig. 2c). Regarding the prediction of whether each position in a pre-mRNA transcript is a splice donor, splice acceptor, or neither, we adapted Nucleotide Transformer model to deliver nucleotide-level splice site predictions and achieved a top-k accuracy of 95% and a precision-recall AUC of 0.98 (Fig. 2d). Notably, our 2.5B 6kb-context model matched the performance of the state-of-the-art SpliceAI-10k [36], which was trained on 15kb input sequences, in addition to other splicing baselines; and outperformed SpliceAI when tested on 6kb input sequences. Finally, in the case of housekeeping and developmental enhancer prediction, our model slightly surpassed (1%) and obtained lower (4%) correlation values respectively (Fig. 2e), when compared to those of DeepSTARR [12]. Across these three different tasks, we also conducted a comparison between our parameter-efficient fine-tuning and full model fine-tuning (i.e. training all the parameters of the whole model to optimize its performance on a specific task or dataset). Interestingly, we observed no significant improvement in chromatin and splicing predictions, and only a modest 3% enhancement in enhancer activity predictions (Supplementary Fig. 2), supporting the use of our efficient fine-tuning approach. Altogether, our extensive benchmarks and results demonstrate the flexibility and performance of Nucleotide Transformer as a general approach to tackle many diverse genomics tasks with high accuracy.

### Benchmark of genomics foundational models

We compared the Nucleotide Transformer models to other genomics foundational models: DNABERT-2 [23], HyenaDNA (1kb and 32kb context length; [25]) and the Enformer (using it as an alternative architecture for a pre-trained model, [19]; see Fig. 2a,b and Methods). We excluded DNABERT-1 from this comparison as it can only handle a maximum input length of 512bp and thus could not be used for most tasks [20]. To ensure fair comparisons, all models underwent fine-tuning and evaluation using the same protocol across the 18 downstream tasks (see Methods). When compared with DNABERT-2, HyenaDNA-32kb and Enformer, our Multispecies 2.5B model achieved the highest overall performance across tasks (Fig. 2a,b, Table 9). Still, Enformer achieved the best performance on enhancer prediction and some chromatin tasks, demonstrating that it can be a strong DNA foundation model. Our model exhibited better performance than every other model on all promoter and splicing tasks. Notably, despite HyenaDNA being pre-trained on the human reference genome, our Multispecies 2.5B model matched (7) or outperformed (11) it in all 18 tasks, highlighting the advantage of pre-training on a diverse set of genome sequences. We have established an interactive leaderboard containing results for all models across each task to facilitate comparisons ^1^. To the best of our knowledge, this represents the most extensive benchmark of foundational genomics models to date and should serve as a reference for the development of further language models in genomics (Fig. 1c).

### The Nucleotide Transformer model learned to reconstruct human genetic variants

To investigate the advantages of increasing the number of parameters in the models and enhancing the genomic diversity of the training datasets, we evaluated the models’ ability to reconstruct masked nucleotides. Specifically, we divided the human reference genome into non-overlapping 6kb sequences, tokenized each sequence into 6-mers, randomly masked a certain number of tokens, and then calculated the proportion of tokens that were accurately reconstructed (Fig. 3a; Supplementary Fig. 4, Methods). We observed that the reconstruction accuracy of the Human reference 500M model exhibited higher median accuracy compared to the 1000G 500M model (median=0.202 versus 0.198, P<2.2e-16, two-sided Wilcoxon rank sum test). However, the accuracy of this model was lower than that achieved by the 1000G 2.5B model (median=0.216; P<2.2e-16, two-sided Wilcoxon rank sum test) and the 2.5B Multispecies model (median=0.219; P<2.2e-16, two-sided Wilcoxon rank sum test) (Supplementary Fig. 4), highlighting the impact of simultaneously increasing the model size and diversifying the dataset. Interestingly, while the Multispecies 2.5B model exhibited the highest overall reconstruction accuracy, specific sequences demonstrated significantly higher reconstruction accuracy for the Human 1000G 2.5B model (Fig. 3b). This likely indicates that certain characteristics of human sequences were better captured and learned by the latter model. For example, it is possible that genomic annotations and molecular phenotypes controlled by elements overlapping these sequences might be better predicted by it.

**Figure 3.**
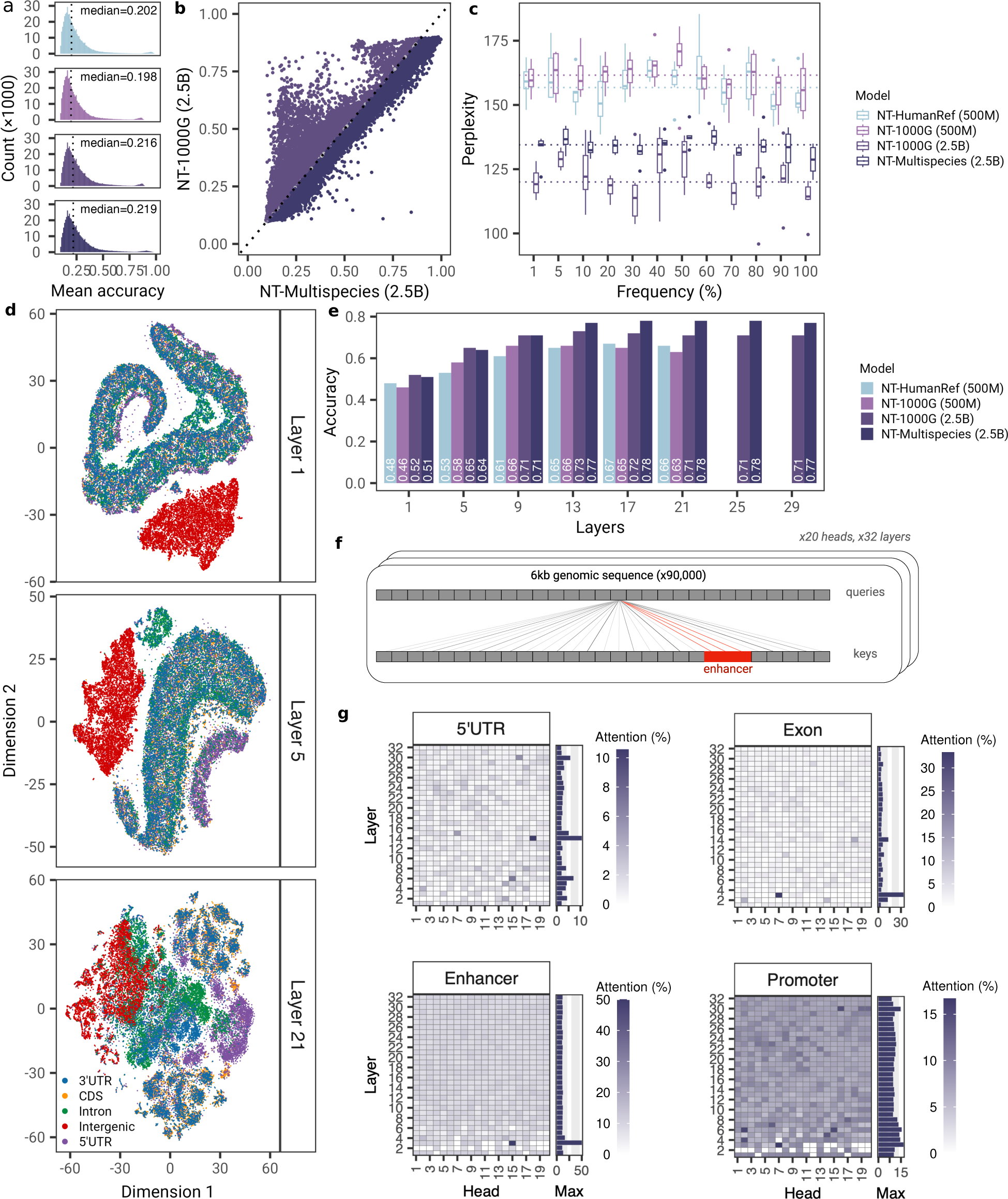
The Nucleotide Transformer models acquired knowledge about genetic variations and genomic elements. **a)** Mean reconstruction accuracy of 6kb sequences across transformer models. **b)** Comparison of reconstruction accuracies between the 1000G 2.5B and Multispecies 2.5B models. **c)** Reconstruction perplexity across allele frequencies. Perplexity values shown per frequency bin are based on 6 meta-populations from the Human Genome Diversity Project. **d)** t-SNE projections of embeddings of 5 genomic elements from layer 1, 5, and 21 based on the Multispecies 2.5B model. **e)** Accuracy estimates based on probing to classify 5 genomic elements across layers. **f)** Schematic describing the evaluation of attention levels at a given genomic element. **g)** Attention percentages per head and layer across the Multispecies 2.5B model computed on 5’ UTR, exon, enhancer, and promoter regions. Barplot on the right of each tile plot shows the maximum attention percentage across all heads for a given layer.

To gain a deeper understanding of how transformer language models represent variant sites in DNA and their capabilities in imputing such variants, we then focused on polymorphisms identified in human samples from diverse populations. In order to assess whether our models can generalize and reconstruct variants in unseen human genome sequences, we evaluated their sequence reconstruction effectiveness by recording model perplexity scores across single nucleotide polymorphisms (SNPs) occurring at frequencies ranging from 1% to 100% (Methods). To account for the influence of genomic background and genetic structure on the mutation reconstruction, we considered an independent dataset of genetically diverse human genomes, originating from 6 different meta-populations [37] (see Methods). Across varying SNP frequencies, we consistently observed that the 2.5B parameter models, with median perplexity across populations ranging from 95.9±17.9 (2SD) to 145±9.3, outperformed their 500M counterparts, for which the perplexity values fluctuate between 138.5 8.4 and 185.4 9.5 (Fig. 3c). Interestingly, the 1000G 2.5B model exhibited lower perplexity scores than the Multispecies 2.5B (median=121.5 versus 134.0, P<1.8e-12, two-sided Wilcoxon rank sum test) indicating that it effectively leveraged human genetic variability observed in the 1000G data to accurately reconstruct variants in unseen human genomes. We confirmed this result by analyzing the models’ variant reconstruction accuracy after excluding variants present in any individual from the 1000 Genomes Project, showing that the 1000G 2.5B model is leveraging relevant genomic features learned across diverse human sequences beyond simply allele frequencies (Supplementary Fig. 5). These results are in line with the observed reconstruction accuracies over human genome sequences, and confirm the impact of model size on performance (Fig. 3b). Notably, and in contrast to typical genotype imputation methods where accuracy declines as variant frequency decreases [38], our models exhibited comparable performance across frequencies, suggesting their potential to enhance genotype imputation even for variants present at lower frequencies.

### The attention layers detect known genomic elements in an unsupervised manner

To gain insights into the interpretability and understand the type of sequence elements that the Nucleotide Transformer models use when making predictions, we explored different aspects of their architecture. First, we assessed the extent to which the embeddings can capture sequence information associated with five genomic elements (Supplementary Table. 10 7). We observed that the Nucleotide Transformer models, without any supervision, learned to distinguish genomic sequences that were uniquely annotated as intergenic, intronic, coding, and UTR regions, albeit with varying degrees of proficiency across different layers (Fig. 3d; Supplementary Fig. 7; Methods). In particular, the 500 million-sized models and those trained on less diverse sequences, exhibited lower separation among genomic regions, reinforcing the enhanced ability of the largest models to capture relevant genomic patterns during self-supervised training. In the case of the Multispecies 2.5B model, the strongest separation at layer 1 was observed between intergenic and non-intergenic regions, followed by 5’ UTR regions on layer 5, and a separation between most regions on layer 21 (Fig. 3d). The limited separation of 3’ UTR regions from other elements suggests that the model has not fully learned to distinguish this type of element, or as previously suggested, that many of these regions might be missannotated [39]. Consistent with these observations, our probing strategy demonstrated high classification performance for these elements, with accuracy values exceeding 0.78, especially for deeper layers (Fig. 3e). This demonstrated that the Nucleotide Transformer models have learned to detect known genomic elements within their embeddings in an unsupervised manner, which can be harnessed for efficient downstream genomics task predictions.

Next, we conducted an analysis of the models through the lens of attention to comprehend which sequence regions are captured and utilized by the attention layers [40]. We computed the attention percentages across each model head and layer for sequences containing nine different types of genomic elements related to gene structure and regulatory features (Fig. 3f). In a formal sense, an attention head is considered to recognize specific elements when its attention percentage significantly exceeds the naturally occurring frequency of that element in the pre-training dataset (Methods). For example, a percentage of 50% implies that, on average over the human genome, 50% of the attention of that particular head is directed toward the type of element of interest. By applying this approach to each type of element across approximately 10,000 different 6kb windows, where the element can be situated at various positions and accounts for between 2-11% of the sequence (Table 10), we discovered that attention is distinctly focused on various types of genomic elements across its diverse heads and layers (Fig. 3g, Supplementary Figs. 8 - 16). The number of significant attention heads across layers varied markedly across models, with the highest number of significant attention heads observed for the Multispecies 2.5B model for introns (117 out of 640 heads), exons (72) and TF binding sites (74) (Supplementary Figs. 8, 9 and Table 12), despite the relatively small proportion of sequences belonging to exons and TF motifs. Regarding enhancers, the maximum attention percentages were highest for the largest models, with the 1000G 2.5B model, for instance, achieving nearly 100% attention (Supplementary Fig. 15). Similar patterns were also observed for other genomic elements such as 3’ UTR, promoter, and TF binding sites, where the 1000G 2.5B model showed highly specialized heads with high attention, particularly in the first layers (Supplementary Figs. 8 - 16).

To gain deeper insights into the pre-trained Nucleotide Transformer Multispecies 2.5B model at higher resolution (i.e. focusing on more local sequence features), we examined token probabilities across different types of genomics elements as a metric of sequence constraints and importance learned by the model. Specifically, we calculated the 6-mer token probabilities (based on masking each token at a time) for every 6kb window in chromosome 22. Our findings revealed that, in addition to repetitive elements, that are well reconstructed by the model as expected, the pre-trained model learned a variety of gene structure and regulatory elements. These included acceptor and donor splicing sites, polyA signals, CTCF binding sites and other genomic elements (Supplementary Fig. 17a-d). Furthermore, we compared our token predictions with an experimental saturation mutagenesis splicing assay of exon 11 of the gene *MST1R* (data from Braun, Simon, et al. [41]). This analysis revealed a significant correlation between the experimental mutation effects and the token predictions made by the Multispecies 2.5B model (Pearson Correlation Coefficient (PCC) = 0.44; Supplementary Fig. 17e). The model not only captured constraints at different splicing junctions but also identified a region in the middle of the second intron crucial for the splicing of this exon. These results serve as a robust validation of the biological knowledge acquired by the Nucleotide Transformer model during unsupervised pre-training.

Lastly, the Multispecies 2.5B model, which had been fully fine-tuned on the DeepSTARR enhancer activity data, was examined to determine whether it had learned about TF motifs and their relative importance specifically for enhancer activity. We used a dataset of experimental mutation of hundreds of individual instances of five different TF motif types across hundreds of enhancer sequences [12] and evaluated the model’s accuracy in predicting these mutation effects. In comparison with the state-of-the-art enhancer activity DeepSTARR model, our model achieved similar performance for four TF motifs and demonstrated superior performance for the Dref motif (Supplementary Fig. 18). Collectively, these results illustrate how the Nucleotide Transformer models have acquired the ability to recover gene structure and functional properties of genomic sequences and integrate them directly into its attention mechanism. This encoded information should prove helpful in evaluating the significance of genetic variants.

### The Nucleotide Transformer embeddings predict the impact of mutations

Additionally, we evaluated the Nucleotide Transformer models’ ability to assess the severity of various genetic variants and prioritize those with functional significance. We first investigated the use of zero-shot scores, which are scores used to predict classes that were not seen by the model during training. Specifically, we computed zero-shot scores using different aspects of vector distances in the embedding space, as well as those derived from the loss function, and compared their distribution across 10 different types of genetic variants that varied in severity [42] (Fig. 4a and Methods). Encouragingly, several of these zero-shot scores exhibited a moderate correlation with severity across the models (Supplementary Fig. 19). This illustrates, how unsupervised training alone captured relevant information related to the potential severity of genetic mutations and highlighted the utility of evaluating different scoring methods. The high variability in correlation between scores also suggests that distinct aspects of the embedding space may more effectively capture information related to severity. Among these scores, cosine similarity exhibited the highest correlation with severity across models, with *r*^2^ values ranging from 0.35 to 0.3 (P-value < 6.55e^−186^) (Supplementary Fig. 19). Across models, the lowest cosine similarity scores were assigned to genetic variants affecting protein function, such as stop-gained variants, as well as synonymous and missense variants (Fig. 4b). Conversely, we noted that higher scores were assigned to potentially less functionally important variants, such as intergenic variants, highlighting its potential use to capture the effect of the severity of genetic variants.

**Figure 4.**
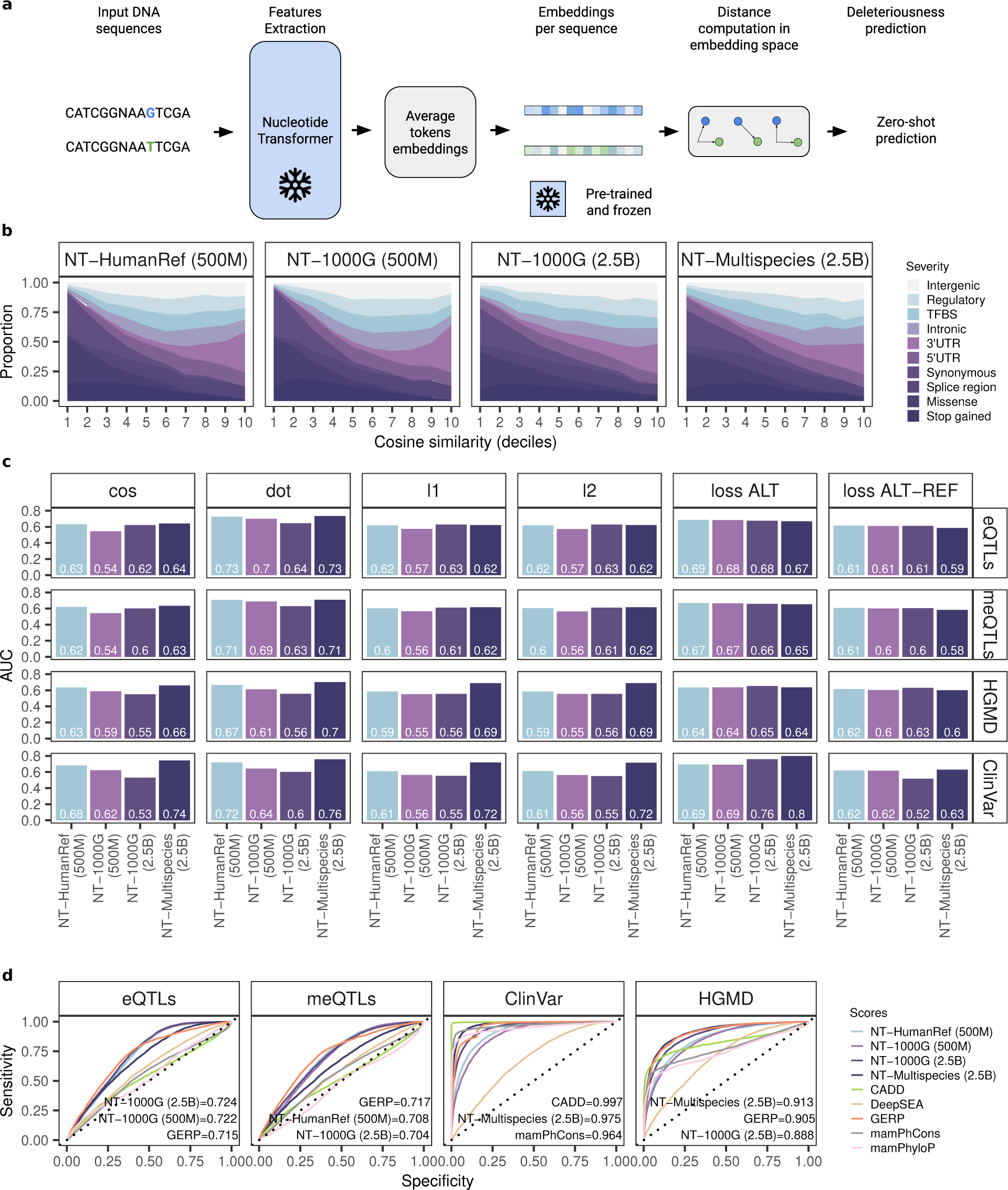
Prioritizing functional genetic variants. **a)** Overview of the Nucleotide Transformer application of zero-shot predictions. **b)** Proportion of variant consequence terms across deciles based on the cosine similarity metric across models. The consequence terms are shown in order of severity (less severe to more severe) as estimated by Ensembl. **c)** Comparison of zero-shot predictions for prioritizing functional variants based on different distance metrics. **d)** Comparison of fine-tuned models and available methods for prioritizing functional variants based on GRASP eQTLs and meQTLs, ClinVar, and HGMD annotated mutations. Model performance is measured with the area under the receiver operating characteristic curve (AUC). The AUC for the three best-performing models is shown.

Next, we also explored the potential of zero-shot scores to prioritize functional variants as well as those with pathogenic effects. Specifically, we assessed the ability of the models to classify genetic variants that influence gene expression regulation (i.e., expression quantitative trait loci [eQTLs]), genetic variants linked to DNA methylation variations (i.e., methylation quantitative trait loci [meQTLs]), genetic variants annotated as pathogenic in the ClinVar database, and genetic variants reported in the Human Gene Mutation Database (HGMD). The zero-shot scores demonstrated high classification performance, with the highest AUCs across the four tasks ranging from 0.7 to 0.8 (Fig. 4c). The highest performance obtained for the Clinvar variants (AUC=0.80 for the Multispecies 2.5B model), suggests that, at least for highly pathogenic variants, zero-shot scores might be readily applicable. Lastly, and to more formally evaluate the effectiveness of these models, we also made predictions based on fine-tuned models and compared their performance against several methods. These methods encompassed those measuring levels of genomic conservation, as well as scores obtained from models trained on functional features. Notably, the fine-tuned models either slightly outperformed or closely matched the performance of the other models (Fig. 4d). The best-performing models for prioritizing molecular phenotypes (i.e. eQTLs and meQTLs) were those trained on human sequences, whereas the best-performing model for prioritizing pathogenic variants was based on multispecies sequences. Given that the most severely pathogenic variants tend to impact gene function due to amino acid changes, it is possible that the multispecies model leveraged sequence variation across species to learn about the degree of conservation across sites. Our results also suggest that higher predictive power for non-coding variants such as eQTLs and meQTLs, could be achieved by better-learned sequence variation derived from increased human genetic variability. Moreover, when compared to zero-shot scores, the dot product yielded AUC values of 0.73 and 0.71 for eQTLs and meQTLs, respectively, slightly surpassing or matching those obtained by the fine-tuned models. Given that most of these genetic variants tend to reside within regulatory regions [43–45], it is probable that the models, without any supervision, have learned to distinguish relevant regulatory genomic features associated with gene expression and methylation variation. This is in accordance with the level of attention observed across layers and heads, especially for relevant regulatory sequences such as enhancers (Fig. 3d) and promoters (Supplementary Fig. 13), which have been shown to be enriched in meQTLs and eQTLs [43–45]. Overall, these results illustrate how DNA-based transformer models can help reveal and contribute to understanding the potential biological implications of variants linked to molecular phenotypes and disease.

### Optimization of Nucleotide Transformer models towards cost-effective predictions in genomics

Finally, we explored the potential for optimizing our top-performing models by incorporating contemporary architectural advancements and extending the training duration. We developed four new Nucleotide Transformer models (NT-v2) with varying number of parameters ranging from 50M to 500M, and introduced a series of architectural enhancements (Supplementary Table 1; Methods). These include the incorporation of rotary embeddings, the implementation of swiGLU activations, and the elimination of MLP biases and dropout mechanisms, in line with the latest research [46]. Additionally, we expanded the context length to cover 12 kbp to accommodate longer sequences and capture more distant genomic interactions. We extended the training duration of the 250 and 500 million parameter models to encompass 1 trillion tokens, aligning with recent recommendations in the literature [47] (Fig. 5a). After pre-training on the same multispecies dataset, all four NT-v2 models underwent fine-tuning and evaluation across the same set of 18 downstream tasks, with their results compared to those of the four initial NT models (Fig. 5b, Supplementary Tables 7 and 9).

**Figure 5.**
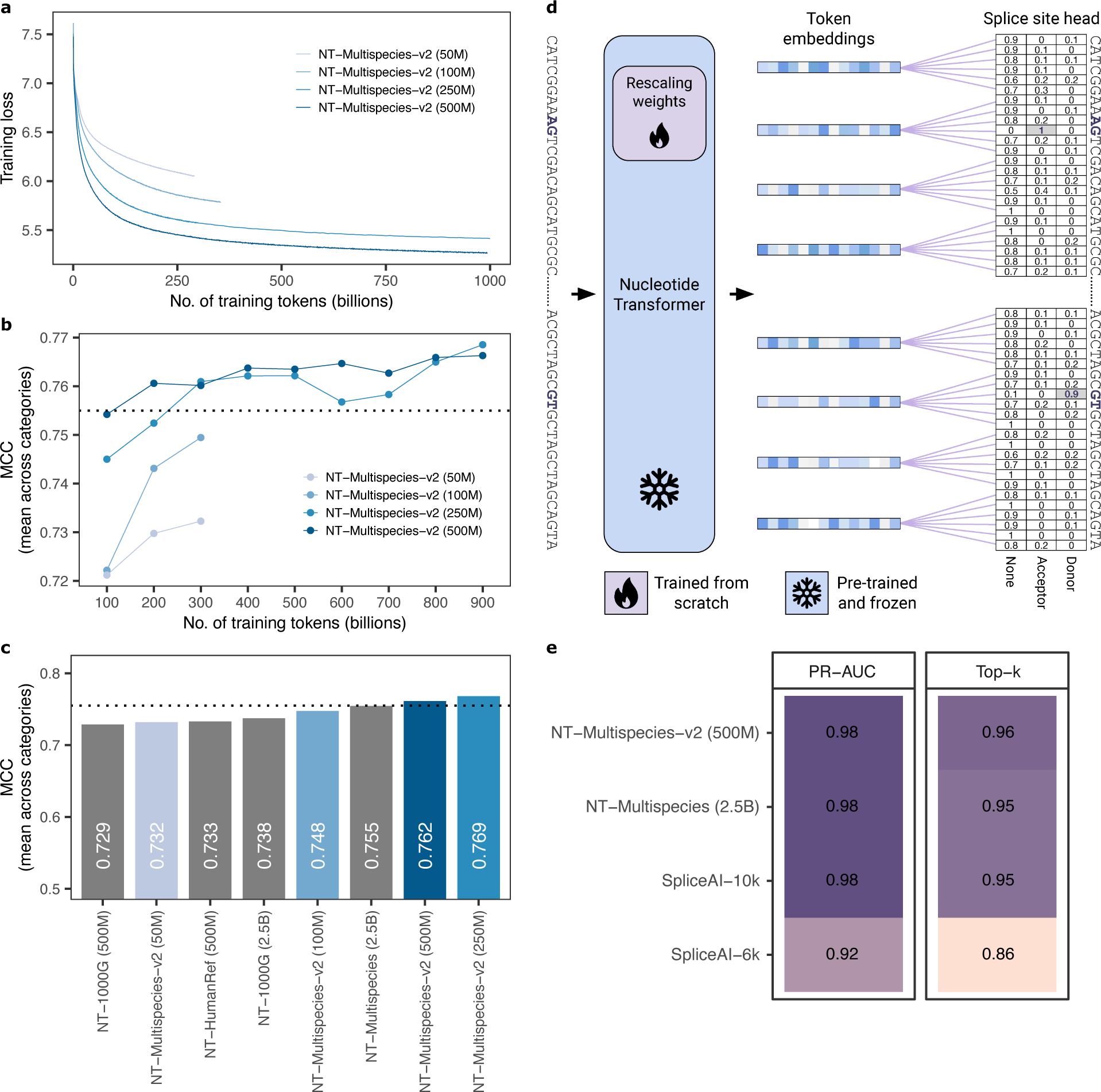
Efficient model architecture allows to match performance while strongly reducing the number of model parameters. **a)** Nucleotide Transformer-v2 models loss value evolution during training as a function of the number of tokens seen so far. **b)** Normalized mean of MCC performance for NT-v2 models as a function of the number of tokens seen during pre-training. **c)** Normalized mean of MCC performance across downstream tasks (divided by category) for all NT (grey) and -v2 (blue shades) Nucleotide Transformer models after fine-tuning. Black dashed line represents the performance of the NT 2.5B multispecies model. **d)** Overview of the Nucleotide Transformer fine-tuning on nucleotide-level splice site prediction task. Pre-trained weights and weights trained from scratch are highlighted. **e)** The Multispecies v2 500M model performance on predicting splice sites from the human genome, compared with its 2.5B counterpart and SpliceAI.

We observed that the 50 million-parameter NT-v2 model achieved similar performance than our two NT 500 million-parameter models as well as the 2.5 billion-parameter model trained on the 1000G dataset. This demonstrates that the synergy of a superior pre-training dataset, coupled with advancements in training techniques and architecture, can lead to a remarkable 50-fold reduction in model parameters while simultaneously enhancing performance (Fig. 1a). Indeed, the NT-v2 250 and 500 million-parameter model managed to achieve improved performance over the 2.5 billion-parameter multi-species model while maintaining a significantly leaner parameter count and doubling the perception field. Of particular interest is the NT-v2 250 million-parameter model, which attained the best performance on our benchmark (average MCC of 0.769) while being 10-times smaller than the 2.5 billion-parameter model (Fig. 5b).

To gain further insights into the need for longer pre-training, we conducted a systematic evaluation of the NT-v2 models’ performance in function of the number of tokens seen during pre-training (Fig. 5c). This showed that the 250 million-parameter model has a small improvement over the 500 million model only after training for 900 billion tokens. In summary, the Nucleotide Transformer-v2 models, equipped with a context length of 12 kbp, are suitable for deployment on cost-effective accelerators due to their compact sizes. Consequently, they offer an economically viable and practical alternative for users seeking to leverage cutting-edge foundational models in their downstream applications.

As a relevant use case, with an established well-performing model, we assessed the advantage of the longer context length of the NT-v2 models by evaluating the 500M model on the SpliceAI splicing task. We evaluated the 500M model as this model showed the highest performance on splicing-related tasks (Supplementary Tables 7 and 9). We adapted our classification head to predict nucleotide-level probabilities of being a splicing donor, acceptor, or none (Fig. 5d; Methods). Compared to the NT 6kb models, our NT-v2 500M 12kb model improved performance by 1% to a top-k accuracy of 96% and a precision-recall AUC of 0.98 (Fig. 5e). This performance surpasses that of the state-of-the-art SpliceAI-10k [36], which was trained on 15kb input sequences. It’s worth noting that we did not attempt to optimize our model architectures specifically for the splicing prediction task; instead we applied a similar fine-tuning approach as used for other downstream tasks, with adjustments in the classification head to yield nucleotide-level predictions. Further architectural refinements tailored to specific tasks like splicing are likely to enhance performance. In summary, these results affirmed the utility and effectiveness of both Nucleotide Transformer v1 and v2 models for a wide range of genomics tasks, requiring minimal modifications and compute power while achieving high accuracy.

## Discussion

To our knowledge, this study represents the first attempt to investigate the impact of different datasets used to pre-train equally-sized transformer models on DNA sequences. Our results, based on distinct genomic prediction tasks, demonstrate that both intra-species (i.e., when training on multiple genomes of a single species) and inter-species (i.e., on genomes across different species) variability significantly influences accuracy across tasks (Fig. 1c, 2a). Models trained on genomes from different species outperform those trained exclusively on human sequences in the considered human prediction tasks. This suggests that transformer models trained on diverse species have learned to capture genomic features that likely have functional importance across species, thus enabling better generalization in various human-based prediction tasks. Based on this finding, we anticipate that future studies may benefit from leveraging genetic variability across species.

The transformer models trained in this study ranged from 50 million up to 2.5 billion parameters, which is twenty times larger than DNABERT-2 [23] and ten times larger than the Enformer [19] backbone models. As previously demonstrated in NLP research [27], our results on the 18 genomic prediction tasks confirmed that increasing model size leads to consistently improved performance. To train the models with the largest parameter sizes, we utilized a total of 128 GPUs across 16 compute nodes for 28 days. Significant investments were made in engineering efficient training routines that fully utilized the infrastructure, underscoring the importance of both specialized infrastructure and dedicated software solutions. Once trained, however, these models can be used for inference at a relatively low cost, and we provide notebooks to apply these models to any downstream task of interest, thereby facilitating further research.

Previous work, which was based on language models trained on biological data (primarily protein sequences), have evaluated downstream performance exclusively by probing the last transformer layer [5]. This choice was likely driven by its perceived ease of use, relatively good performance, and low computational complexity. In this study, our aim was to evaluate downstream accuracy through computationally intensive and thorough probing of different transformer layers, downstream models, and hyperparameter sweeps. We observed that the best probing performance was achieved with intermediate transformer layers (Supplementary Fig. 1), which aligns with recent work in computational biology [34] and common practice in NLP [48]. Through probing alone, the Multispecies 2.5B model outperformed the BPNet baseline model for 8 out of 18 tasks. Additionally, we explored a recent downstream fine-tuning technique that introduces a small number of trainable weights into the transformer. This approach provides a relatively fast and resource-efficient fine-tuning procedure with minor differences compared to full-model fine-tuning (IA^3^ [35]). Notably, this fine-tuning approach only requires 0.1% of the total number of parameters, allowing even our largest models to be fine-tuned in under 15 minutes on a single GPU. In comparison to the extensive probing exercise, this technique yielded superior results while using fewer compute resources, confirming that downstream model engineering can lead to performance improvements [49]. The use of this technique makes fine-tuning competitive with probing from an operational perspective, both for training and inference, while achieving improved performance.

While supervised models with optimized architectures and extensive training on large datasets continue to demonstrate superior performance, the DNA language approach remains competitive with these models in a diverse range of tasks encompassing areas such as histone modifications, splicing sites, and regulatory elements characterization. Moreover, the Nucleotide Transformer model represents a versatile approach that can be seamlessly tailored to a diverse array of tasks for both human and non-human species. The value of our unsupervised pre-training approach becomes particularly evident when dealing with smaller datasets, where training supervised models from scratch typically yields limited results. Recognizing the pivotal role of foundational genomics models in the field, we have conducted an extensive comparison and benchmarking study, evaluating our model against four distinct pre-trained models with different architectures: DNABERT-2 [23], HyenaDNA [25] and Enformer [19]. This robust benchmark will serve as a reference point for the development of future language models in genomics.

Through various analyses related to the transformer architecture, we have demonstrated that the models have acquired the ability to recognize key regulatory genomic elements. This property is demonstrated throughout the analysis of attention maps, embedding spaces, token reconstruction, and probability distributions. Crucial regulatory elements governing gene expression, such as enhancers and promoters [43–45], were consistently detected by all models across multiple heads and layers. Additionally, we observed that each model contained at least one layer that produced embeddings clearly distinguishing the five genomic elements analyzed. Given that self-supervised training facilitated the detection of these elements, we anticipate that this approach can be harnessed for the characterisation or discovery of novel genomic elements in future research.

We have demonstrated that transformer models can match and even outperform other methods for predicting variant effects and deleteriousness. In addition to developing supervised transformer models, we have also showcased the utility of zero-shot-based scores, especially for predicting non-coding variant effects. Given that these zero-shot-based scores can be derived solely from genomic sequences, we encourage their application in non-human organisms, especially those with limited functional annotation.

Lastly, we have demonstrated the potential of enhancing the model architecture to achieve a dual benefit of reducing model size and improving performance. Our subsequent series of NT-v2 models have a context length of 12 kbp. To put this in perspective, this length is 24x and 4x larger than the respective average context lengths of DNABERT-2 (3kb). These advanced models not only exhibit improved downstream performance but also offer the advantage of being suitable for execution and fine-tuning on economical hardware. This advantage arises from their compact size, complemented by the utilization of the efficient fine-tuning techniques that we have introduced.

While our models have an attention span that remains limited, the recently developed Enformer model [19] suggested that increasing the perception field up to 200kb is necessary to capture long-range dependencies in the human genome. The authors argued that these are needed to accurately predict gene expression, which is controlled by distal regulatory elements, usually located far greater than 10-20kb away from the transcription starting site. Processing such large inputs is intractable with the standard transformer architecture, due to the quadratic scaling of self-attention with respect to the sequence length. The Enformer model addresses this issue by passing sequences through convolution layers to reduce input dimensions before reaching the transformer layers. However, this choice hampers its effectiveness in language modeling. On the other hand, the recent HyenaDNA models [25] were trained with perception fields up to 1M base pairs, but our benchmark analyses in downstream tasks show that the performance of these models quickly deteriorates when the perception field used during training increases. Based on our results, we suggest that developing transformer models with the ability to handle long inputs while maintaining high performance on shorter ones is a promising direction for the field. In an era where multi-omics data are rapidly expanding, we ultimately anticipate that the methodology presented here, along with the available benchmarks and code, will stimulate the adoption, development, and enhancement of large foundational language models in genomics.

## Acknowledgements

We thank members of the Rostlab, particularly Tobias Olenyi, Ivan Koludarov, and Burkhard Rost for constructive discussions that helped identify interesting research directions. We also thank Maša Roller for helpful discussion and commenting on the manuscript. Furthermore, we would like to express our gratitude to all volunteers who generously provided their biological data, as well as to the researchers who deposited experimental data in public databases, the researchers who maintain these databases, and those who make analytical and predictive methods available to the scientific community.

## Methods

### Models

Language models (LMs) have been primarily developed within Natural Language Processing (NLP) to model spoken languages [1, 2]. A LM is a probability distribution over sequences of tokens (often words), i.e. given any sequence of words, an LM will return the probability for that sentence to exist. LMs gained in popularity thanks to their ability to leverage large unlabeled datasets to generate general-purpose representations that can solve downstream tasks even when little supervised data is available [50]. One technique to train LMs tasks models to predict the most likely tokens at masked positions in a sequence, often referred to as masked language modelling (MLM). Motivated by results obtained with MLM in the field of protein research [4, 5], where proteins are considered as sentences and amino-acids as words, we apply MLM to train language models transformers in genomics, considering sequences of nucleotides as sentences and k-mers (with k=6) as words. Transformers are a class of deep learning models that achieved breakthroughs in machine learning fields including NLP and computer vision. They consist of an initial embedding layer that transform positions in the input sequence into an embedding vector, followed by stack of self-attention layers that sequentially refine these embedding. The main technique to train language models transformers with MLM is called Bidirectional Encoder Representations from Transformers (BERT) [1]. In BERT, all positions in the sequence can attend to each other allowing the information to flow in both directions, which is essential in the context of DNA sequences. During training, the final embedding of the network is fed to a language model head that transforms it into a probability distribution over the input sequence.

### Architecture

All our models follow an encoder-only transformer architecture. An embedding layer transforms sequences of tokens into sequences of embeddings. Positional encodings are then added to each embedding in the sequence to provide the model with positional information. We use a learnable positional encoding layer that accepts a maximum of 1000 tokens. We used 6-mer tokens as a trade-off between sequence length (up to 6kb) and embedding size, and because it achieved the highest performance when compared with other token lengths. The token embeddings are then processed by a transformer layer stack. Each transformer layer transforms its input through a layer normalisation layer followed by a multi-head self-attention layer. The output of the self-attention layer is summed with the transformer layer input through a skip connection. The result of this operation is then passed through a new layer normalisation layer and a two-layer perceptron with GELU activations [51]. The number of heads, the embedding dimension, the number of neurons within the perceptron hidden layer and the total number of layers for each model can be found in Table 1. During self-supervised training, the embeddings returned by the final layer of the stack are transformed by a language model head into a probability distribution over the existing tokens at each position in the sequence.

Our second version Nucleotide Transformer v2 models include a series of architectural changes that proved more efficient: instead of using learned positional embeddings, we use Rotary Embeddings [52] that are used at each attention layer; we use Gated Linear Units with swish activations without bias, making NLPs more efficient. These improved models also accept sequences up to 2, 048 tokens leading to a longer context window of 12 kbp.

### Training

The models are trained following the BERT methodology [1]. At each training step a batch of tokenized sequences is sampled. The batch size is adapted to available hardware and model size. We conducted all experiments on clusters of A100 GPUs, and took batches of sizes 14 and 2 sequences to train the 500M and 2.5B parameters models, respectively. Within a sequence, of a subset of 15% of tokens, 80% are replaced by a special mask [MASK] token. For training runs on the Human reference genome and multispecies datasets, an additional 10% of the 15% subset of tokens are replaced by randomly selected standard tokens (i.e. any token different from the class [CLS], pad [PAD] or mask [MASK] token), as was done in BERT. For training runs on the 1000G dataset, we skipped this additional data augmentation, as the added noise was greater than the natural mutation frequency present in the human genome. For each batch, the loss function was computed as the sum of the cross-entropy losses, between the predicted probabilities over tokens and the ground truth tokens, at each selected position. Gradients were accumulated to reach an effective batch size of 1M tokens per batch. We used the Adam optimizer [53] with a learning rate schedule, and standard values for exponential decay rates and epsilon constants, β_1_ = 0.9, β_2_ = 0.999 and ϵ=1e-8. During a first warmup period, the learning rate was increased linearly between 5e-5 and 1e-4 over 16k steps before decreasing following a square root decay until the end of training.

We slightly modified the hyperparameters of our NT-v2 models: the optimizer and learning rate schedule are kept the same, however we increased batch size to 512 (1,000,000 tokens per batch). Inspired by Chinchilla scaling laws [47], we also trained our NT-v2 models for longer duration compared to other deep learning models. Specifically, we pre-trained our NT-v2 50M and 250M parameters models for 300B tokens, while our 250M and 500M parameters models were trained for up to 1 trillion tokens to understand the scaling laws at play. In comparison, the NT-v1 2.5B parameters models were trained for 300B tokens, and their 500M counterparts were trained for 50B tokens. In the end we used the following model checkpoints for the NT-v2 models: checkpoint 300B tokens for 50M and 100M models, checkpoint 800B tokens for the 250M model, and checkpoint 900B tokens for the 500M model.

### Probing

We refer to probing the assessment of the quality of the model embeddings to solve downstream tasks. After training, for each task, we probe each layer of the model and compare several downstream methods to evaluate in depth the representations capabilities of the model. In other words, given a dataset of nucleotide sequences for a downstream task, we compute and store the embeddings returned by ten layers of the model. Then, using the embeddings of each individual layer as inputs, we trained several downstream models to solve the downstream task. We tested logistic regression with the default hyperparameters from scikit-learn [54] and a multi-layer peceptron. As we observed that the choice of hyperparameters, such as the learning rate, the activation function and the number of layers per hidden layer impacted final performance, we also ran hyperparameters sweeps for each downstream model. We used a 10-fold validation scheme, where the training dataset was split ten times in a training and validation set, that contain different shuffles with 90% and 10% of the initial set. For a given set of hyperparameters, ten models were trained over the ten splits, and their validation performances were averaged. This procedure is run 100 times with a Tree-structured Parzen Estimator solver [55] guiding the search over the hyperparameters space, before evaluating the best performing set of models on the test set. Therefore, for each downstream task, for ten layers of each pre-trained model, the performance on the test set is recorded at the end of the hyperparameters search. The hyperparameters of the best performing probe across the pre-trained models and their layers are reported in Table 8. This probing strategy resulted in 760,000 downstream models trained, which provides detailed analysis into various aspects of training and using LMs, such as the role of different layers on downstream task performance.

As a baseline, we evaluated the performance of a logistic regression model that takes as input the tokenized sequence, i.e. before passing the tokens through the transformer layers. Using the raw tokenized sequences as input yielded much better performance than using a vector where the token ids were one-hot encoded and passed through a pooling layer (summing or averaging, over the sequence length axis).

### Fine-tuning

In addition to probing our models through embedding extraction at various layers, we also performed parameter-efficient fine-tuning through the IA^3^ technique [35]. Using this strategy, the language model head is replaced by either a classification or regression head depending on the task at hand. The weights of the transformer layers and embedding layers are frozen and new, learnable weights are introduced. For each transformer layer, we introduced three learned vectors *l*_*k*_ ∈ R^*d*^_*k*_, *l*_*v*_ ∈ R^*d*^_*v*_ and *l*_ff_ ∈ R^*d*^_ff_, which were introduced in the self-attention mechanism as:

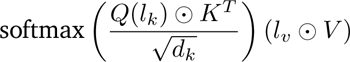

and in the position-wise feed-forward networks as (*l*_ff_γ (*W*_1_*x*)) *W*_2_, where γ is the feed-forward network nonlinearity, and ⨀ represents the element-wise multiplication. This adds a total of *L* (*d*_*k*_ + *d*_*v*_ + *d*_ff_) new parameters, where *L* is the number of transformer layers. We refer to these learnable weights as *rescaling* weights. The intuition is that during fine-tuning these weights will weigh the transformer layers to improve the final representation of the model on a downstream task, so that the classification/regression head can more accurately solve the problem. As we observed layer specialization during probing, we speculate that this fine-tuning technique will similarly select layers with greater predictive ability for particular tasks.

In practice, the number of additional parameters introduced by rescaling weights and the classification/regression head weights represented approximately 0.1% of the total number of weights of the model. This increased fine-tuning speed since just a fraction of parameters needed updating. Similarly, it alleviated storage requirements, needing to create space for just 0.1% new parameters over 500M and 2.5B for each downstream task using traditional fine-tuning. For instance, for the 2.5B parameters models, the weights represent 9.5GB. Considering 18 downstream tasks, classical fine-tuning would have required 9.5 × 18 = 171 GBs, whereas parameter-efficient fine tuning required only 171 MB.

Like in the probing scheme, the training dataset was split ten times in a training and validation set, that contain different shuffles with 90% and 10% of the initial set. For each, split, the model was fine-tuned for 10k steps and parameters yielding the highest validation score were then used to evaluate the model on the test set. We used a batch size of eight and the Adam optimizer with a learning rate of 3e-3. Other optimizer parameters were maintained from training regiments. Each model is fine-tuned for 10k steps for each task. These hyperparameters were selected as they led to promising results in the field of NLP [35]. Diverging hyperparameter choices did not yield significant gains in our experiments. We have also compared this approach with fine-tuning from a randomly initialized checkpoint.

### Comparison with supervised BPNet baseline

As baseline supervised models, we trained different variants of the BPNet convolutional architecture [9] from scratch on each of the 18 tasks. We tested the original architecture (121 thousand parameters), a large one (28 million) and an extra-large (113 million). For each, the hyperparameters were manually adjusted to yield best performance on the validation set. We implemented a 10-fold cross-validation strategy to measure the performance of each of the 18 models per architecture.

### Comparison with published pre-trained genomics models

We compared the fine-tuned performance of Nucleotide Transformer models on the 18 downstream tasks with three different pre-trained models: DNABERT-2 [23], HyenaDNA (1kb and 32kb context length; [25]) and Enformer [19]. We excluded DNABERT-1 from this comparison as it can only handle a maximum input length of 512bp and thus could not be used for most tasks [20]. We ported the architecture and trained weights of each model to our code framework and performed parameter-efficient fine-tuning on the transformer part of every model as described above, using the same cross-validation scheme for a fair comparison. All results can be visualized in an interactive leader-board ^2^. Only for HyenaDNA we performed full fine-tuning due to the incompatibility of our parameter-efficient fine-tuning approach with the model architecture.

Note that the Enformer has been originally trained in a supervised fashion to solve chromatin and gene expression tasks. For the sake of benchmarking, we re-used the provided model torso as a pre-trained model for our benchmark, which is not the intended and recommended use of the original paper. Though we think this comparison is interesting to highlight the differences between self-supervised and supervised learning for pre-training and observe that the Enformer is a very competitive baseline even for tasks that differ from gene expression.

### Pre-training datasets

#### The Human reference genome dataset

The Human reference dataset was constructed by considering all autosomal and sex chromosomes sequences from reference assembly GRCh38/hg38 ^3^ and reached a total of 3.2 billion nucleotides.

#### The 1000G dataset

To inform the model on naturally occurring genetic diversity in humans, we constructed a training dataset including genetic variants arising from different human populations. Specifically, we downloaded the variant calling format (VCF) files^4^ from the 1000 Genomes project [56], which aims at recording genetic variants occurring at a frequency of at least 1% in the human population. The dataset contained 3,202 high-coverage human genomes, originating from 27 geographically structured populations of African, American, East Asian, and European ancestry as detailed in Table 2, making up a total of 20.5 trillion nucleotides. Such diversity allowed the dataset to encode a better representation of human genetic variation. To allow haplotype reconstruction in the FASTA format from the VCF files, we considered the phased version of the data, which corresponded to a total of 125M mutations, 111M and 14M of which are single nucleotide polymorphisms (SNPs) and indels, respectively.

#### The Multispecies dataset

The selection of genomes for the multispecies dataset was primarily based on two factors: (1) the quality of available reference genomes and (2) diversity among the species used. The genomes chosen for this dataset were selected from the ones available at the Reference Sequence (RefSeq) collection of NCBI^5^. To ensure a diverse set of genomes, we randomly selected one genome at the genus level from each of the main groupings available in RefSeq (i.e., archaea, fungi, vertebrate_mammalian, vertebrate_other, etc.). However, due to the large number of available bacterial genomes, we opted to include only a random subset of them. Plant and virus genomes were not taken into account, as their regulatory elements differ from those of interest in this work. The resulting collection of genomes was downsampled to a total of 850 species, whose genomes add up to 174 billion nucleotides. The final contribution of each class, in terms of number of nucleotides, to the total number of nucleotides in the dataset, displayed in Table 3, is the same as in the original collection parsed from NCBI. Finally, we enriched this dataset by selecting several genomes that have been heavily studied in the literature (Table 4).

### Data preparation

Once the FASTA files of each genome / individual were collected, they were assembled into one unique FASTA file per dataset that was then pre-processed before training. During this data processing phase, all nucleotides other than A,T,C,G were replaced by N. A tokenizer was employed to convert strings of letters to sequences of tokens. The tokenizer used as alphabet the 4^6^ = 4096 possible 6-mer combinations obtained by combining A,T,C,G, as well as five extra tokens to represent stand-alone A,T,C,G and N. It also included three special tokens, namely the padding [pad], masking [mask] and the beginning of sequence (also called class; [CLS]) token. This adds to a vocabulary of 4104 tokens. To tokenize an input sequence, the tokenizer will start with a class token and then convert the sequence starting from the left, matching 6-mer tokens when possible, or falling back on the stand-alone tokens when needed (for instance when the letter N is present or if the sequence length is not a multiple of 6).

For the multispecies and Human reference dataset, genomes are split into overlapping chunks of 6100 nucleotides, each sharing the first and last 50 nucleotides with the previous and last chunk, respectively. As a data augmentation exercise, for each epoch and chunk, a starting nucleotide index is randomly sampled between 0 and 100, and the sequence is then tokenized from this nucleotide until 1000 tokens is reached. The number of epochs was determined depending on the dataset so that the model processed a total of 300B tokens during training. At each step, a batch of sequences sampled randomly within the epoch set was fed to the model. For the 1000G dataset, batches of sequences from the Human reference genome, prepared as specified above, are sampled at each step. Then, for each sampled chunk, an individual from the 1000G dataset is randomly selected, and if that individual carries mutations at the positions and chromosome corresponding to that chunk, these mutations are introduced into the sequence, and the corresponding tokens replaced. This data processing technique ensured uniform sampling both over the genome and over the individuals during training, as well as enabled to efficiently store only mutations for each individual, instead of full genomes.

#### Hardware

All models were trained on the Cambridge-1 Nvidia supercomputer system, using 16 nodes, each equipped with eight A100 GPUs, leading to a total of 128 A100 GPUs used. During training, model weights were replicated on each GPU, while batches were sharded across GPUs. Gradients were computed on each shard and accumulated before being averaged across devices and backpropagated. We relied on the jax library^6^ that relied on the NCCL^7^ protocol to handle communications between nodes and devices, and observed almost linear decrease of the training time with respect to the number of GPU available. The 500M parameters models were trained on a single node for a day, while the 2.5B models models required the whole cluster for 28 days to be trained. The Nucleotide Transformer version 2 (NT-v2) models with varying number of parameters ranging from 50M to 500M were similarly trained on a single node for a single day. All fine-tuning runs were performed on a single node with eight A100 GPUs. As for the training runs, the models weights were replicated and batches distributed across GPUs. As we used a batch size of eight for fine-tuning, each GPU processed a single sample before averaging the gradients and applying them. On average, a fine-tuning run lasted 20 minutes for the 500M parameter models, and 50 minutes for the 2.5B parameter models.

For the probing experiments, all embeddings (for all sequences in all downstream tasks, for selected layers of each model) were computed and stored on a single node with eight A100 GPUs, requiring two days to compute. Then, 760,000 downstream models were fit on a cluster of 3000 CPUs, requiring 2.5 days.

### Nucleotide Transformer Downstream tasks

#### Epigenetic marks prediction

Histone ChIP-seq data for 10 histone marks in the K562 human cell line were obtained from ENCODE [31] (https://www.encodeproject.org/). We downloaded bed narrowPeak files with the following identifiers: H3K4me3 (ENCFF706WUF), H3K27ac (ENCFF544LXB), H3K27me3 (ENCFF323WOT), H3K4me1 (ENCFF135ZLM), H3K36me3 (ENCFF561OUZ), H3K9me3 (ENCFF963GZJ), H3K9ac (ENCFF891CHI), H3K4me2 (ENCFF749KLQ), H4K20me1 (ENCFF909RKY), H2AFZ (ENCFF213OTI). For each dataset, we selected 1kb genomic sequences containing peaks as positive examples and all 1kb sequences not overlapping peaks as negative examples.

#### Promoter sequence prediction

We built a dataset of promoter sequences to evaluate the capabilities of the model to identify promoter motifs. We downloaded all human promoters from the Eukaryotic Promoter Database [30] (https://epd.expasy.org/epd/), spanning 49bp upstream and 10bp downstream of transcription start sites (file: https://epd.expasy.org/ftp/epdnew/H_sapiens/006/Hs_EPDnew_006_hg38.bed). This resulted in 29,598 promoter regions, 3,065 of which were TATA-box promoters (using the motif annotation at https://epd.expasy.org/ftp/epdnew/H_sapiens/006/db/promoter_motifs.txt). We selected 300bp genomic sequences containing promoters as positive examples and all 300bp sequences not overlapping promoters as negative examples. These positive and negative examples were used to create three different binary classification tasks: presence of any promoter element (Promoter all), a promoter with a TATA-box motif (Promoter TATA) or a promoter without a TATA-box motif (Promoter non-TATA).

#### Enhancer sequence prediction

Human enhancer elements were retrieved from ENCODE’s SCREEN database (https://screen.wenglab.org/) [32]. Distal and proximal enhancers were combined. Enhancers were split in tissue-specific and tissue-invariant based on the vocabulary from Meuleman et al. [33] https://www.meuleman.org/research/dhsindex/. Enhancers overlapping regions classified as tissue-invariant were defined as that, while all other enhancers were defined as tissue-specific. We selected 400bp genomic sequences containing enhancers as positive examples and all 400bp sequences not overlapping enhancers as negative examples. We created a binary classification task for the presence of enhancer elements in the sequence (Enhancer) and a multi-label prediction task with labels being tissue-specific enhancer, tissue-invariant enhancer or none (Enhancer types).

#### Splice site prediction

We obtained all human annotated splice sites from GENCODE [29] V44 gene annotation. Annotations were filtered to exclude level 3 transcripts (automated annotation), so all training data was annotated by a human. We used *extract_splice_sites.py* from HISAT2 [57] (https://github.com/DaehwanKimLab/hisat2/blob/master/hisat2_extract_splice_sites.py) to extract respective splice site annotations. We selected 600bp genomic sequences containing a splice acceptor or donor site in the center as positive examples and all 600bp sequences not overlapping splice sites as negative examples. We used these sequences to create three different tasks to evaluate splice prediction: a multi-label prediction task with labels being acceptor, donor or none (Splice site all); a binary classification task for the prediction of splice acceptors (Splice acceptor); and a binary classification task for the prediction of splice donors (Splice donor).

#### Dataset splits and performance evaluation

Model training and performance evaluation were performed on different sets of chromosomes from the human genome. Namely, sequences from the chromosomes 20 and 21 are used for test and the remaining are used for training the different models. Sequences with Ns were removed. We balanced each dataset by subsampling the negative examples to the same number of positive examples. To obtain small-sized datasets that can be used to benchmark any new design choice or model quickly, we further randomly subsampled examples to a maximum of 30,000 for training and 3,000 for validation and testing (see details in Supplementary Table 5).

We used a 10-fold cross-validation scheme to evaluate each model, where the training set is split in 10 folds and each time we use 9 of those parts for training and reserve one tenth for validation and selecting the final checkpoint. We repeat this procedure 10 times per model, each time reserving a different tenth for validation and evaluating the final performance on the same held-out test set. We used the median performance across the 10 folds as the final performance metric of each model on a given task.

For a consistent performance evaluation across tasks, we used the binary or multi-class Matthews Correlation Coefficient (MCC) as metric as it is robust to uneven label ratios. For a final comparison across models, we calculated for each model the mean MCC across the 3 different categories of task (chromatin profiles, regulatory elements and splicing), where for each category we use the median MCC across tasks.

### Additional Downstream tasks

#### Chromatin profiles prediction

We used the DeepSEA dataset^8^ compiled in Zhou et al. 2015 [10] for chromatin profiles prediction. The dataset is composed of 2.4 million sequences, each of size 1000 nucleotides, and associated with 919 chromatin features. These include 690 transcription factor (TF), 125 DNAse, and 104 histone features. As in the original publication, our model is trained simultaneously on the 919 classification tasks, with 919 independent classification heads, and a loss taken as the average of the cross entropy losses. Since each label is highly unbalanced and is composed mostly of negative samples, the losses associated with positive samples are upscaled by a factor of 8. Contrarily to the DeepSEA method [10], which trained two models independently, one on the forward sequences and one on the corresponding reverse-complementary, and evaluated the average of their predictions, the model presented here was trained only on the forward sequences.

#### SpliceAI benchmark

We used the scripts available at the Illumina Basespace platform ^9^ to reproduce the training dataset presented in SpliceAI [36]. Briefly, this training dataset is constructed using GENCODE v24lift37 annotations and RNA-seq data from the GTEx cohort, focusing solely on splice site annotations from the principal transcript. The training dataset comprises annotations from genes located on chromosomes 2, 4, 6, 8, and 10-22, as well as chromosomes X and Y, while annotations from genes on the remaining chromosomes, which were not paralogs, constitute the test dataset. Each sequence produced by this pipeline has a length of 15,000 bp, with the central 5,000 bp containing the sites to be predicted. Additional details about the construction of the training dataset can be found in the original publication. We adapted this original SpliceAI dataset to be able to run our models by reducing the sequence length to 6,000 bp (for NT-v1 model) and 12,000 bp (for NT-v2 model), reducing the flanking contexts but keeping the central 5,000 bp. We also removed sequences that contained Ns. When comparing against SpliceAI on the first dataset, which we refer to as SpliceAI-6k, we appended 9,000 “N” nucleotides as flanking sequence since SpliceAI is based on a model with a 15,000 bp input. When comparing against SpliceAI on the second dataset, we report the performance presented in the original publication, which, compared to this dataset, includes a sequence length of 15,000 bp instead of 12,000 bp.

This task is a multi-label classification for each of the input sequence’s nucleotides similar to Splice AI [36] (Fig. 5d). From each embedding outputted by the transformer model, a head predicts, for each of the 6 nucleotides represented by the token embedding, three label probabilities: splice acceptor, splice donor or none. The head is a simple classification layer that predicts 18 classes, i.e 3 labels for each of the 6 nucleotides. To ensure that each embedding is associated with a 6-mer, the sequences are cut so that their length is divisible by 6. Furthermore, all sequences with Ns are removed from both training and test set, which represents a negligible portion of the data. Note that if we were to use a Byte Pair Encoding tokenizer like DNABERT-2 [23], the number of nucleotides represented by each embedding would vary and make nucleotide-level prediction tasks substantially trickier to implement.

#### Enhancer activity prediction

We used the DeepSTARR enhancer activity dataset ^10^ released in de Almeida et al. 2022 [12]. The dataset is composed of 484,052 DNA sequences of size 249 nucleotides, each measured for their quantitative enhancer activity towards a developmental or a housekeeping promoter. We added two independent regression heads to our models to predict both enhancer activities in simultaneous. Following the methodology used in [3], we chose to treat this regression task as a multi-label classification problem. Specifically, each label *y* was discretized over a set of 50 values (*b*_*i*_)_*i*∈[1,50]_, evenly spaced between the minimum and maximum value. For each label, the model predicts the normalized weights (*w*_*i*_)_*i*∈[1,50]_ such that

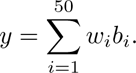

### Additional Performance Analysis

#### t-SNE projections of embeddings

T-distributed Stochastic Neighbor Embedding (t-SNE) was used to reduce Nucleotide Transformer inner embeddings to 2-D vectors to visualize the separation of different genomic elements. Nucleotide Transformer embeddings for each genomic element were computed at several transformer layer outputs and the mean embeddings computed across the sequence locations corresponding to the element were calculated. These mean embeddings were then passed as input into a TSNE reducer object with default parameters from the sklearn python package [58].

### Reconstruction accuracy and perplexity

We studied how pre-trained models could reconstruct masked tokens. We considered a trained language model with parameters *θ*. Within a nucleotide sequence **s** of interest, we masked tokens using one of two strategies (i.e. we replaced the tokens at these positions with the mask token [MASK]). We either masked the central token of the sequence only, or we masked randomly 15% of the tokens within the sequence. The masked sequence is then fed to the model and the probabilities over tokens at each masked position are retrieved. The loss function *l* (*θ*, **s**) and the accuracy *acc*(*θ*, **s**) are defined as follows:

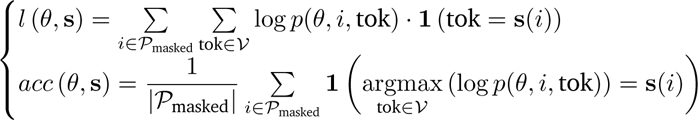

where *P*_masked_ is the set of masked positions and *V* is the vocabulary, i.e. the set of all existing tokens. The perplexity is usually defined in the context of autoregressive generative models. Here, we rely on an alternative definition used in Rives [4], and define it as the exponential of the loss function computed over the masked positions:

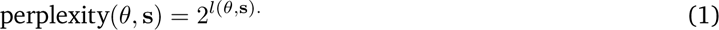

Perplexity measures how well a model can reconstruct masked positions, and is a more refined measure than accuracy, as it does also account for magnitude. In contrast to accuracy, lower perplexity suggests better reconstruction ability, thus better performance.

### Reconstruction of tokens in different genomics elements

We also performed this token reconstruction approach across the full chromosome 22 in 6kb windows. We only kept windows without Ns. For each window masked each token at a time, recovering the predicted probability for the original token in the sequence. We display these scores as a WashU Epigenome Browser session ^11^. To obtain average token probabilities across different types of genomic elements, we retrieved gene annotation regions from Ensembl (exons, introns, splice acceptors and donors, 5’UTR, 3’UTR, TSS, TES ^12^), polyA signal sites from GENCODE ^13^ and regulatory elements from ENCODE (enhancers, promoters and CTCF-bound sites from SCREEN database ^14^).

### Functional variant prioritization

To obtain genetic variants with varying levels of associated severity we used the Variant Effect Prediction (VEP) software [42] and annotated sequences across the human genome. Specifically, we randomly sampled sequences throughout the human genome, and kept genetic variants within those sequences annotated to any of the following classes: “intron variant”, “intergenic variant”, “regulatory region variant”, “missense variant”, “3 prime UTR variant”, “synonymous variant”, “TF binding site variant”, “5 prime UTR variant”, “splice region variant”, and “stop gained variant”. After keeping only single nucleotide polymorphisms and filtering out variants annotated to more than one consequence (e.g. those annotated as stop gained and splice variants), we obtained a final dataset composed of 920 genetic variants per class.

As a positive set of functional genetic variants we compiled SNPs from four different resources. We used SNPs associated to gene expression (i.e. expression quantitative trait loci [eQTLs]) and to methylation variation (i.e., meQTLs) from the Genome-Wide Repository of Associations Between SNPs and Phenotypes (GRASP) database [59] with a P-value <10^−12^, SNPs with “likely pathogenic” annotations from ClinVar [60] and SNPs reported in The Human Gene Mutation Database (HGMD) (public version 2020.4) [61]. After these filters, we retained a total of 80,590, 11,734, 70,224, and 14,626 genetic variants for the eQTLs, meQTLs, ClinVar, and HGMD SNP datasets, respectively. For each of these four datasets, we then constructed a set of negative variants based on SNPs from the 1000 Genomes Project with a minor allele frequency (MAF) >5%, that did not overlap with any variant reported in the dataset tested, and that were within 100kb of the associated variants, resulting in four balanced datasets.

To compute zero-shot based scores for a given site of interest we did the following: For each SNP, we obtained a 6,000 bp sequence centered on the SNP of interest based on the human reference genome. We then created two sequences one carrying the reference allele and a second carrying the alternative allele at the SNP position. We then computed several zero-shot scores that capture different aspects of the vector distances in the embedding space between those two sequences, namely: the L1 distance (Manhattan), (ii) the L2 distance (Euclidean), (iii) the cosine similarity, and (iv) the dot-product (not normalized cosine similarity). We also computed the loss of the alternative allele and the difference in the loss between the sequence carrying the alternative and reference alleles, as two additional zero-shot scores. In the case of functional variants, in addition to zero-shot scores, we also fine-tuned the transformer models to classify positive and negative variants. We employed a similar strategy as the one previously described, with the primary difference being that the training and test sets were divided by chromosomes and strictly kept non-overlapping. Specifically, we divided the 22 chromosomes into 5 sets and sequentially used each of them as a test set and the 4 others as a training set. By fine-tuning on the training set, we could derive probabilities of being a positive variant for each sequence in the test set. We use those probabilities as a score for each SNP.

To compare these predictions against other methods, we randomly sampled 10,000 positive and negative SNPs from each of the four datasets. We then used the Combined Annotation Dependent Depletion tool (version GRCh38-v1.6) to compute CADD, GERP, phastCons, and phyloP scores. The DeepSEA scores were computed using the Beluga model available at: https://hb.flatironinstitute.org/sei/. The score considered was the “disease impact score”reported for each SNP.

### Attention Maps Analysis

We analysed how attention maps gathered from the pre-trained models capture key genomic elements. We followed a methodology proposed in previous work [40]. For a genomic element, we define the indicator function over tokens *f*(*i*) that equals 1 if one or several nucleotides within token *i* belong to the element, and 0 otherwise. We computed the average proportion of attention focused on that genomic element, in one attention head, aggregated over a dataset of nucleotide sequences **X** as:

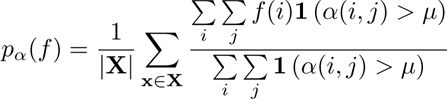

where *α*(*i*, *j*) is the attention coefficient between tokens *i* and tokens *j* defined such that 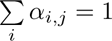 and *μ* is a confidence threshold.

We computed the values of *p*_*α*_(*f*) for all the heads and all the layers of all models, and considered nine elements (“5’ UTR”, “3’ UTR”, “exon”, “intron”, “enhancer”, “promoter”, “CTCF binding site”, “open chromatin”, and “transcription factor binding sites”). We perform these analyses over a dataset made of 90,000 sequences, 10,000 per feature, of length 6k bp extracted from the Human reference genome. The average proportion of tokens belonging to each element can be found in Table 10. For each sequence, the position of the feature within the sequence was sampled uniformly during the dataset creation. As suggested in previous work [40], we selected a confidence threshold *μ* = 0.3 for all experiments.

We considered that a feature is captured by an attention head if the quantity *p*_*α*_(*f*) is significantly greater than the natural occurring frequency of the feature within the dataset (Table 10). To validate this, we conducted a two proportion z-test with the null hypothesis as the natural frequency of the feature, and the alternate hypothesis as *p*_*α*_(*f*). The total number of heads of each model is used as a Bonferroni correction to the significance level, *α*, of 0.05. We computed z-scores and associated p-value for each head in every model for every genomic element as follows:

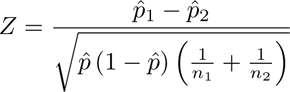

where 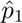 represents the proportion of attention above *μ* associated with each genomic element, *p*^_2_ represents the proportion of the sequence occupied by the genomic element, *n*_1_ is the total number of sequence positions with attention above *μ*, *n*_2_ is the total number of sequence positions. Attention heads with p-values below the Bonferonni corrected significance level are considered to be significant.

### Prediction of important TF motif instances from DeepSTARR data

We retrieved experimental mutagenesis data from the DeepSTARR dataset [12] where individual TF motif instances are mutated and their impact is measured in the activity of developmental and housekeeping enhancers. We assessed the performance of the fully-finetuned NT 2.5B multispecies model (since it was the model with the highest test set performance) to predict the contribution of each TF motif instance by predicting the activity of the wildtype and respective motif-mutant sequence and calculating their log2 fold-change. We compared our predicted mutation effects with the ones predicted by the original method DeepSTARR and the experimentally derived log2 fold-changes.

## Data availability

The Nucleotide Transformer pre-training sequences were obtained from publicly available resources. The 1000 Genomes Project sequences were obtained from http://ftp.1000genomes.ebi.ac.uk/vol1/ftp/data_collections/1000G_2504_high_coverage, and the human and multispecies reference genomes from https://ftp.ncbi.nlm.nih.gov/genomes/refseq/. Gene annotations were obtained from GENCODE (https://www.gencodegenes.org/) and Ensembl databases (https://www.ensembl.org). Variants effects predictions were obtained using the Variant Effect Prediction (VEP) API from Ensembl (https://www.ensembl.org/info/docs/tools/vep/index.html). Pathogenic and regulatory variants were extracted from ClinVar (https://ftp.ncbi.nlm.nih.gov/pub/clinvar/vcf_GRCh38/), the Genome-Wide Repository of Associations Between SNPs and Phenotypes (GRASP) (https://grasp.nhlbi.nih.gov), and The Human Gene Mutation Database (HGMD) (public version 2020.4) through Ensembl Biomart. The Nucleotide Transformer downstream tasks were curated from the following publicly available resources: ENCODE [31–33] (https://www.encodeproject.org/, https://screen.wenglab.org/, https://www.meuleman.org/research/dhsindex/), the Eukaryotic Promoter Database [30] (https://epd.expasy. org/epd/) and GENCODE [29] (see Supplementary Table 5). We have also created an interactive browser session with the pre-trained model token probabilities across the full chromosome 22 on the WashU Epigenome Browser at https://shorturl.at/jov28. HuggingFace versions of our pre-training and downstream tasks datasets can be found at https://huggingface.co/InstaDeepAI.

## Code availability

Model code and weights of the pre-trained transformer models as well as inference code in Jax are available for research purposes at https://github.com/instadeepai/nucleotide-transformer. HuggingFace versions of the models, in PyTorch, can be found at https://huggingface.co/InstaDeepAI. Example note-books are available on HuggingFace at https://huggingface.co/docs/transformers/notebooks#pytorch-bio.

## Supplementary Figures

**Supplementary Figure 1.**
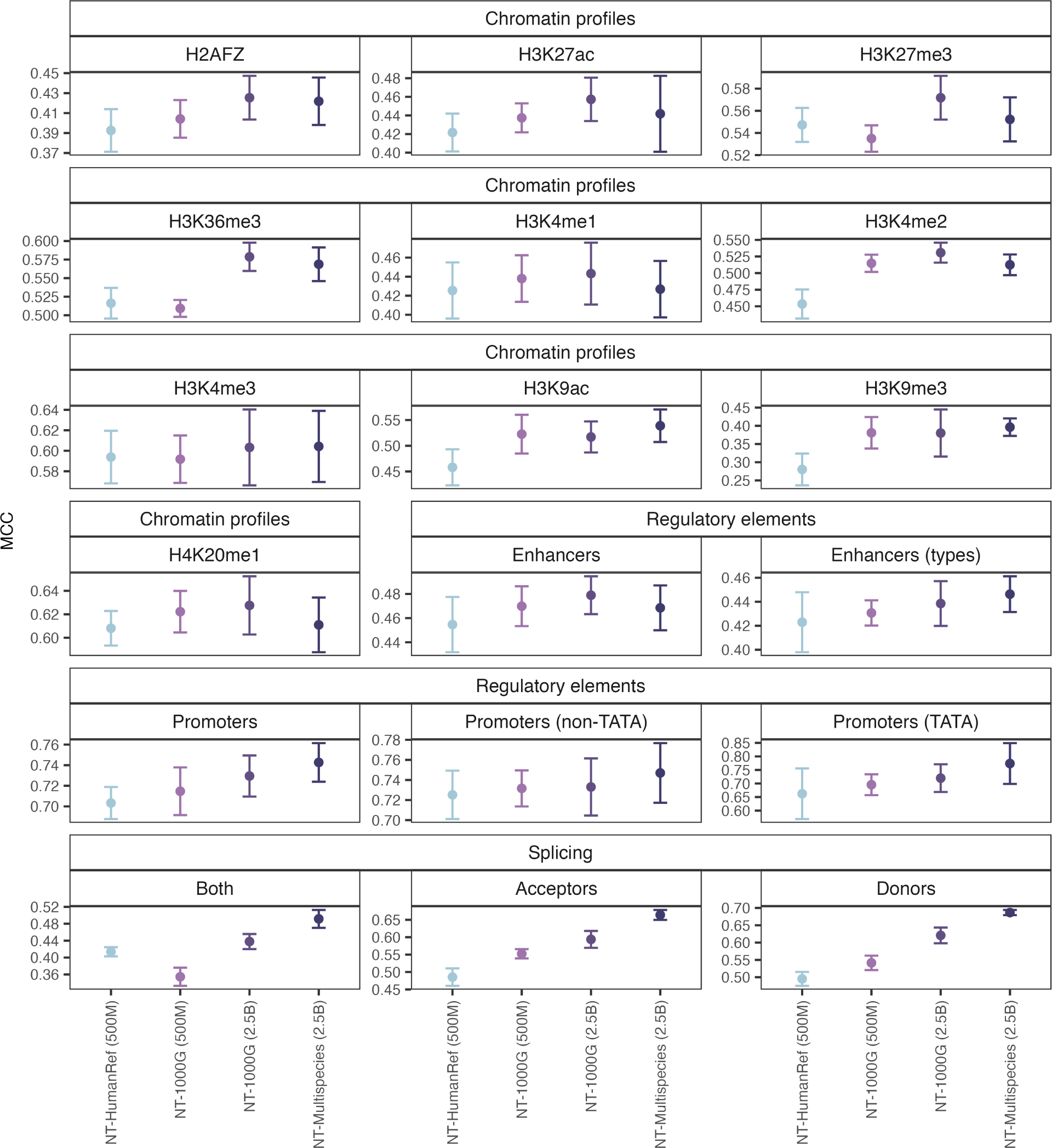
Probing performance results on test sets across downstream tasks for each Nucleotide Transformer model. The performances of the best layers and best downstream models are shown. The error bars represent 2 standard deviations from the 10-fold cross-validation procedure.

**Supplementary Figure 2.**
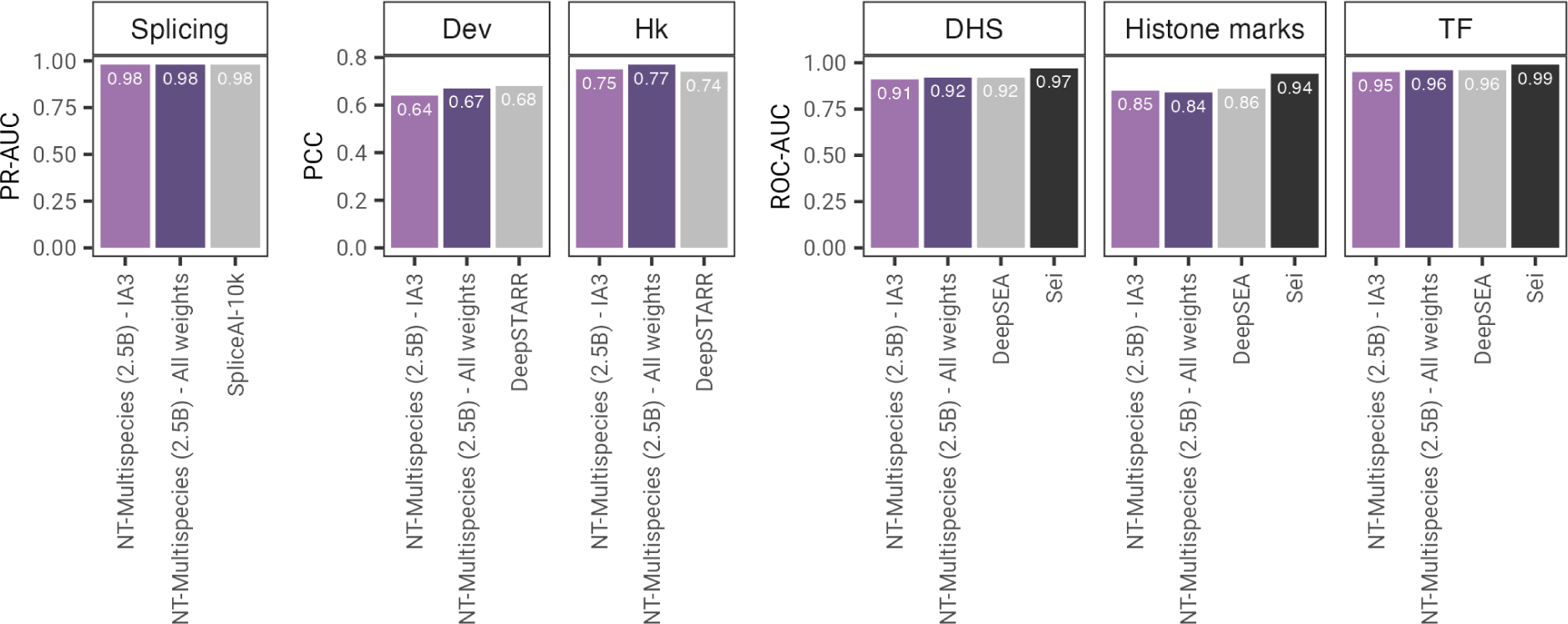
Parameter efficient fine-tuning compared with full-model fine-tuning. The performances across different tasks are shown and compared with respective baselines.

**Supplementary Figure 3.**
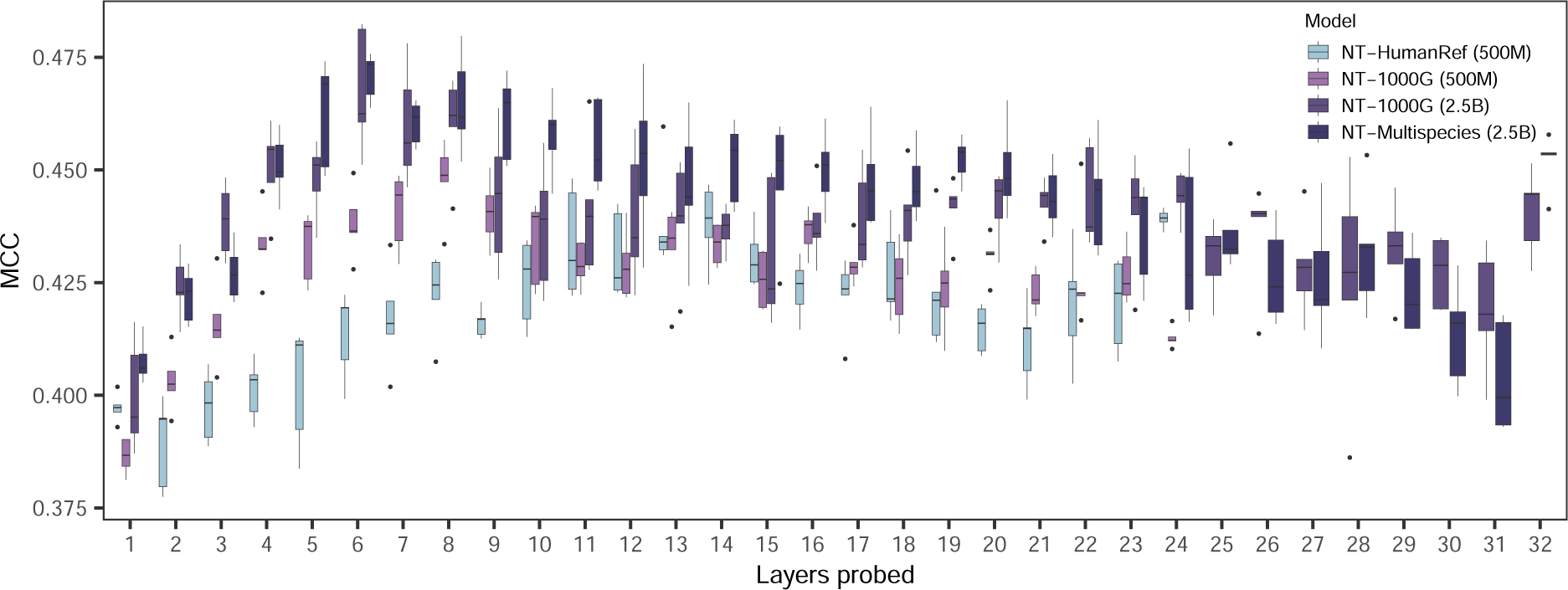
Probing performance across layers for the enhancer (types) prediction task. Boxplots show the MCC values across layers based on 10 fold cross-validation experiments.

**Supplementary Figure 4.**
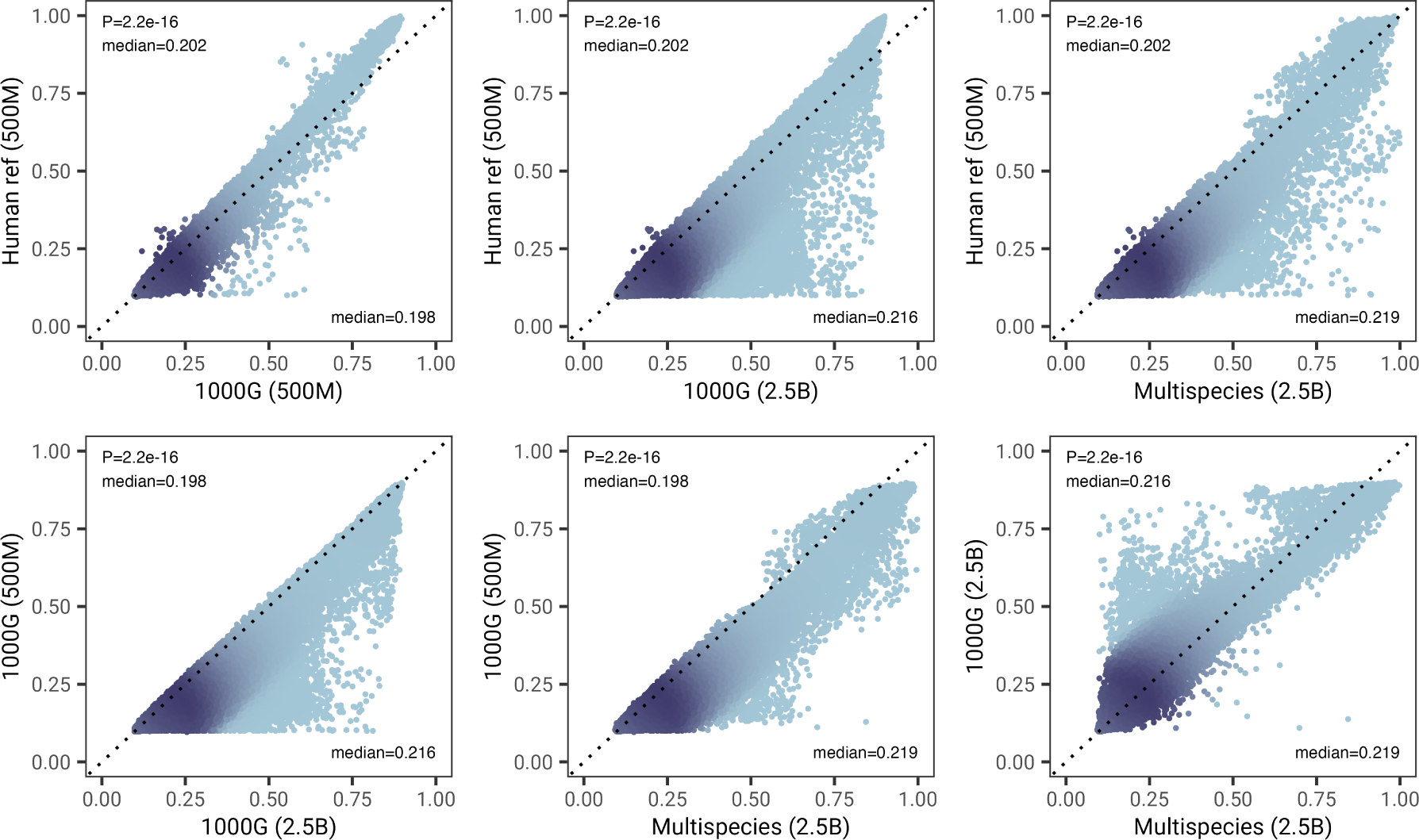
Pairwise comparison of token reconstruction accuracy across Nucleotide Transformer models. P-values refer to a two-sided Wilcoxon signed rank test between models. Median values for the two models compared are shown.

**Supplementary Figure 5.**
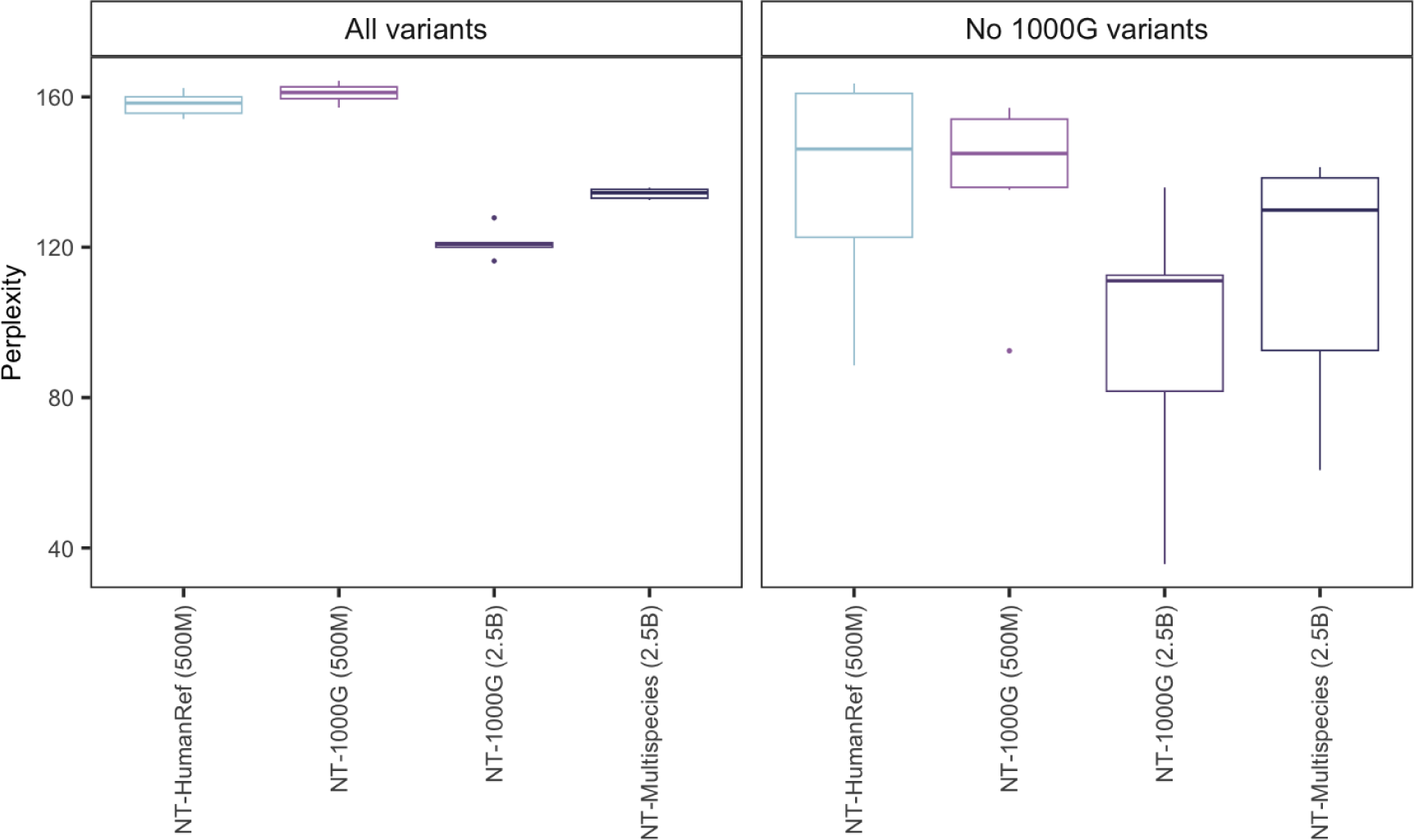
Reconstruction perplexity per model for variants present in 6 meta-populations from the Human Genome Diversity Project. Individual data points in the boxplots refer to the median perplexity values across all variants for each meta-population. Results are shown for all variants or for variants that are not present in the 1000G samples used for model pre-training.

**Supplementary Figure 6.**
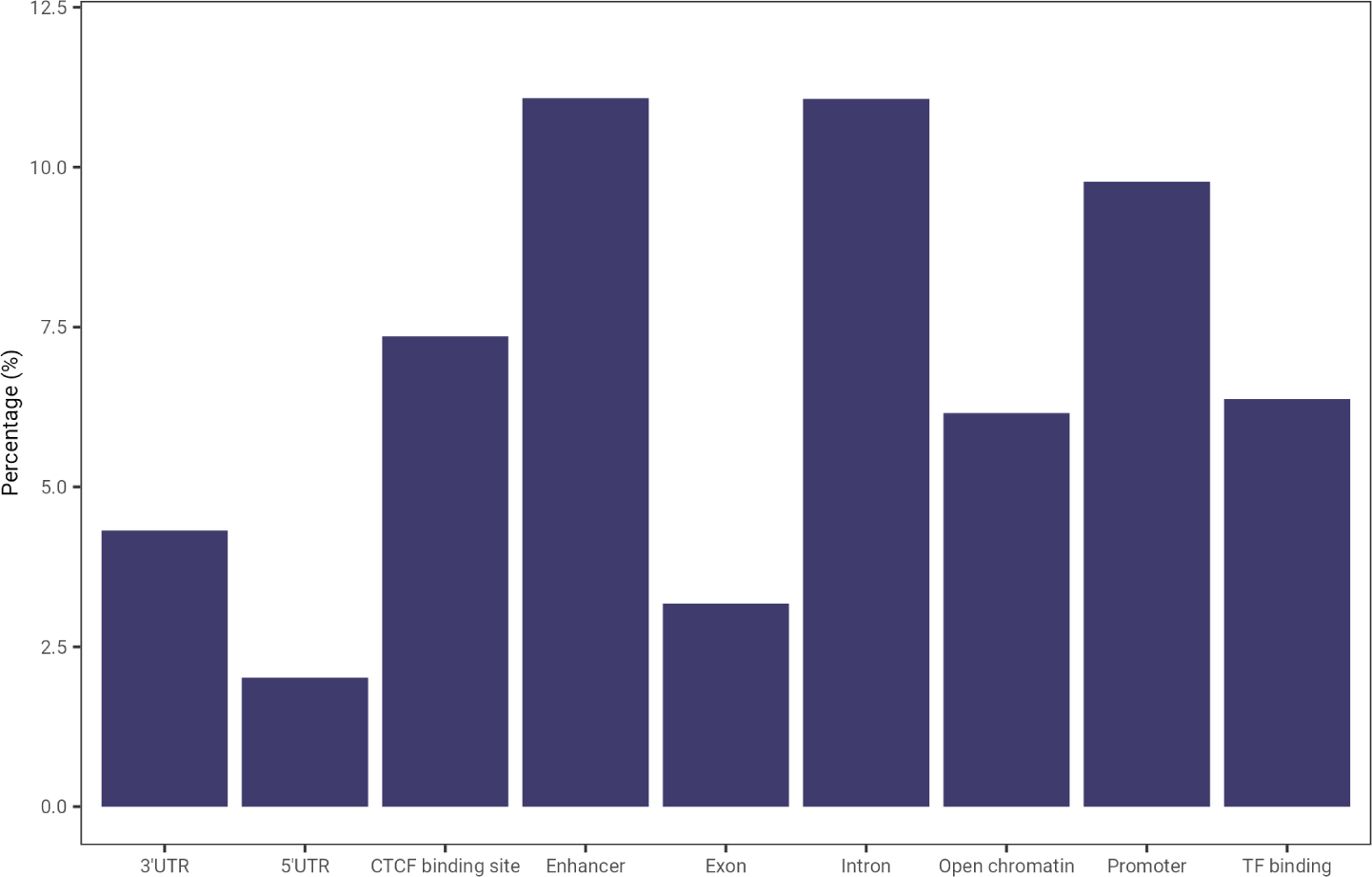
Genomic element length’s percentage of input sequence (6kbp). This corresponds to the dataset used for attention maps and embedding spaces visualization experiments. Details in Table 10.

**Supplementary Figure 7.**
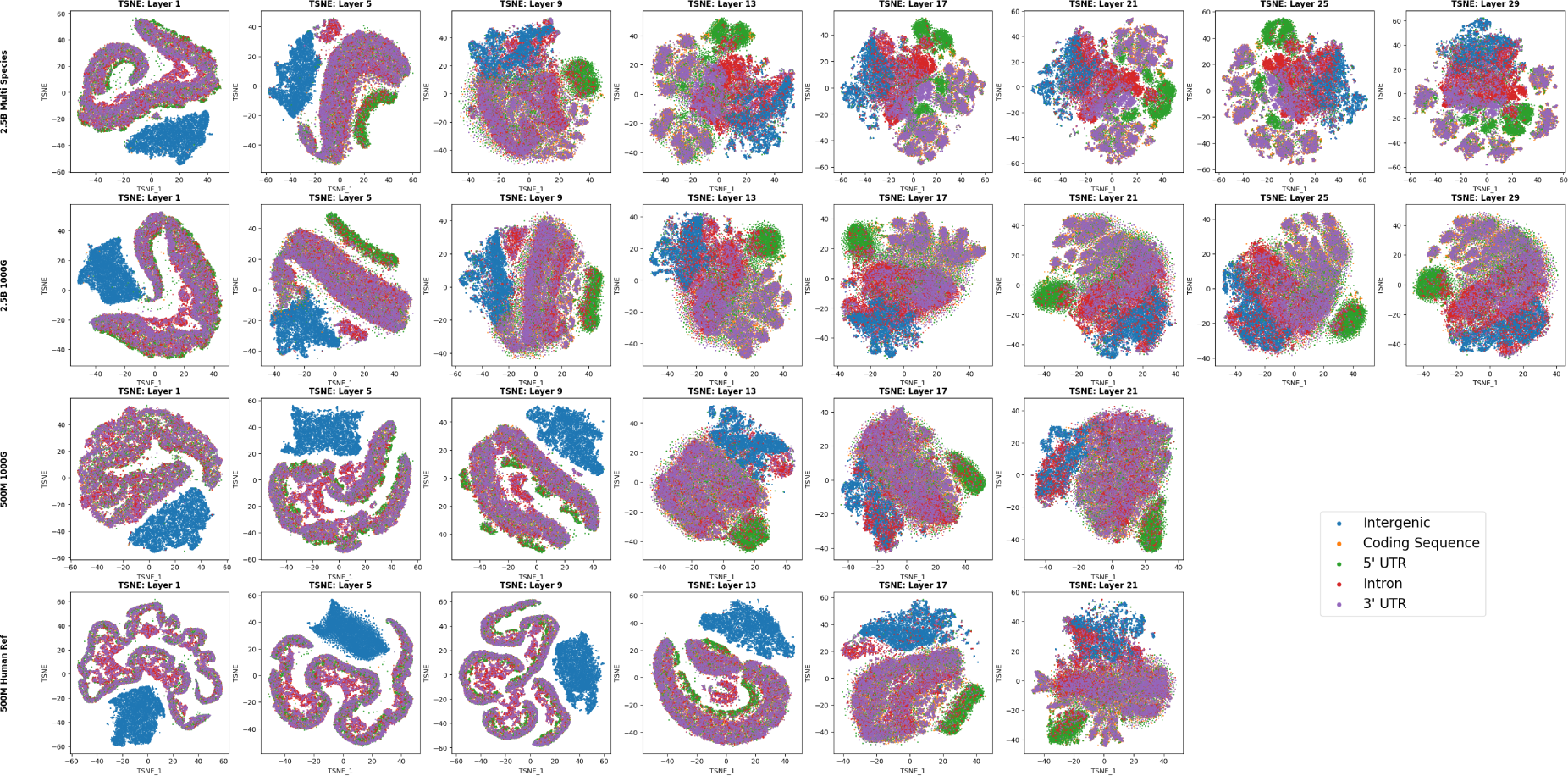
T-SNE projections of embeddings of 5 genomic elements across transformer models and layers.

**Supplementary Figure 8.**
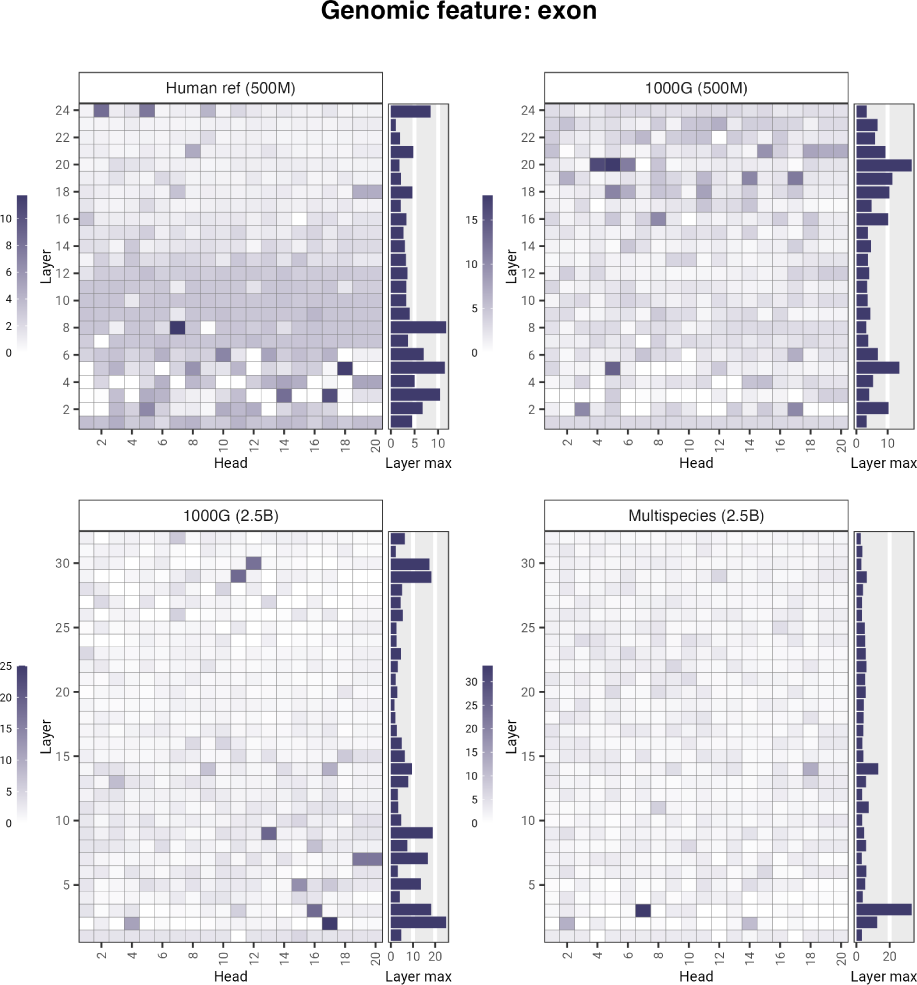
Nucleotide Transformer models’ attention percentages per head computed for exons.

**Supplementary Figure 9.**
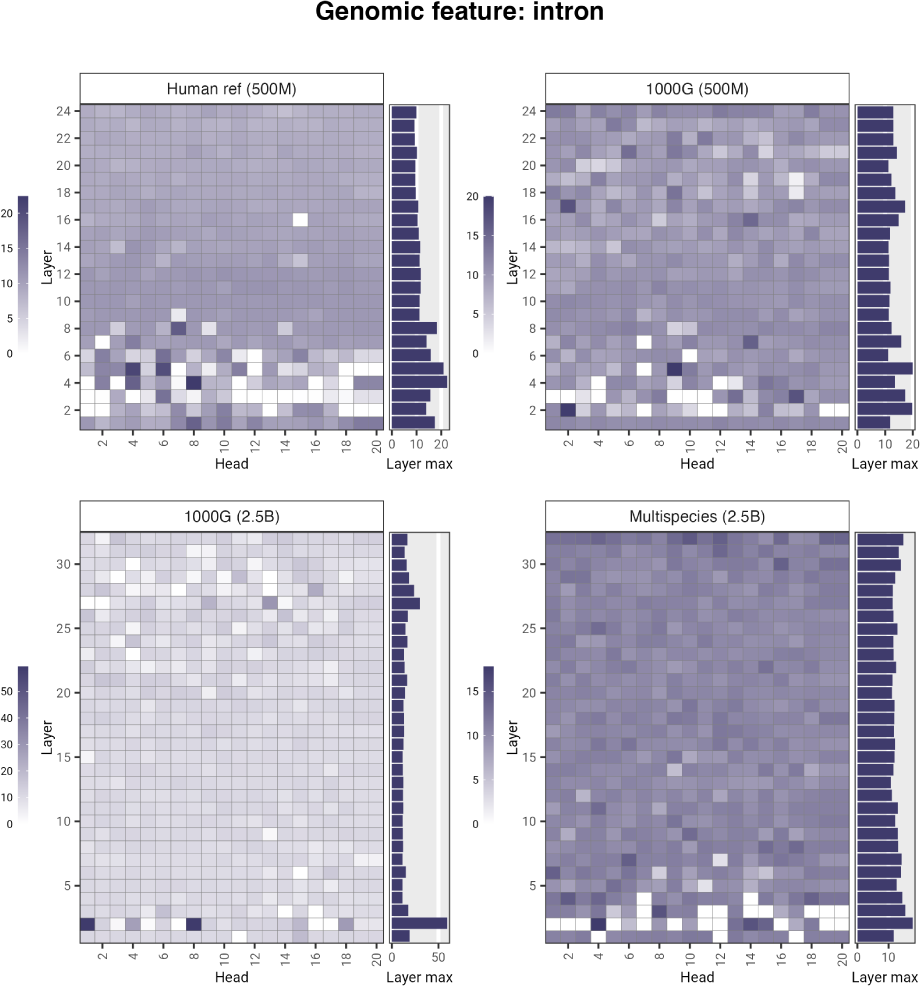
Nucleotide Transformer models’ attention percentages per head computed for introns.

**Supplementary Figure 10.**
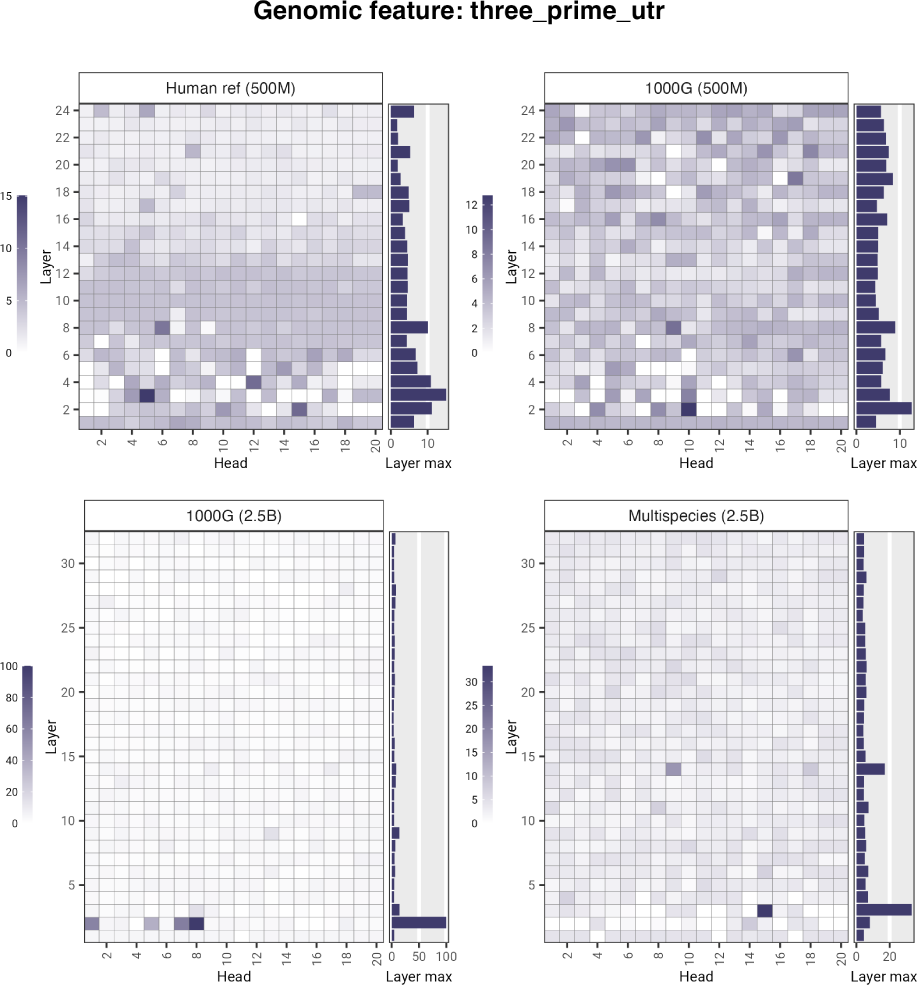
Nucleotide Transformer models’ attention percentages per head computed for 3’ UTR regions.

**Supplementary Figure 11.**
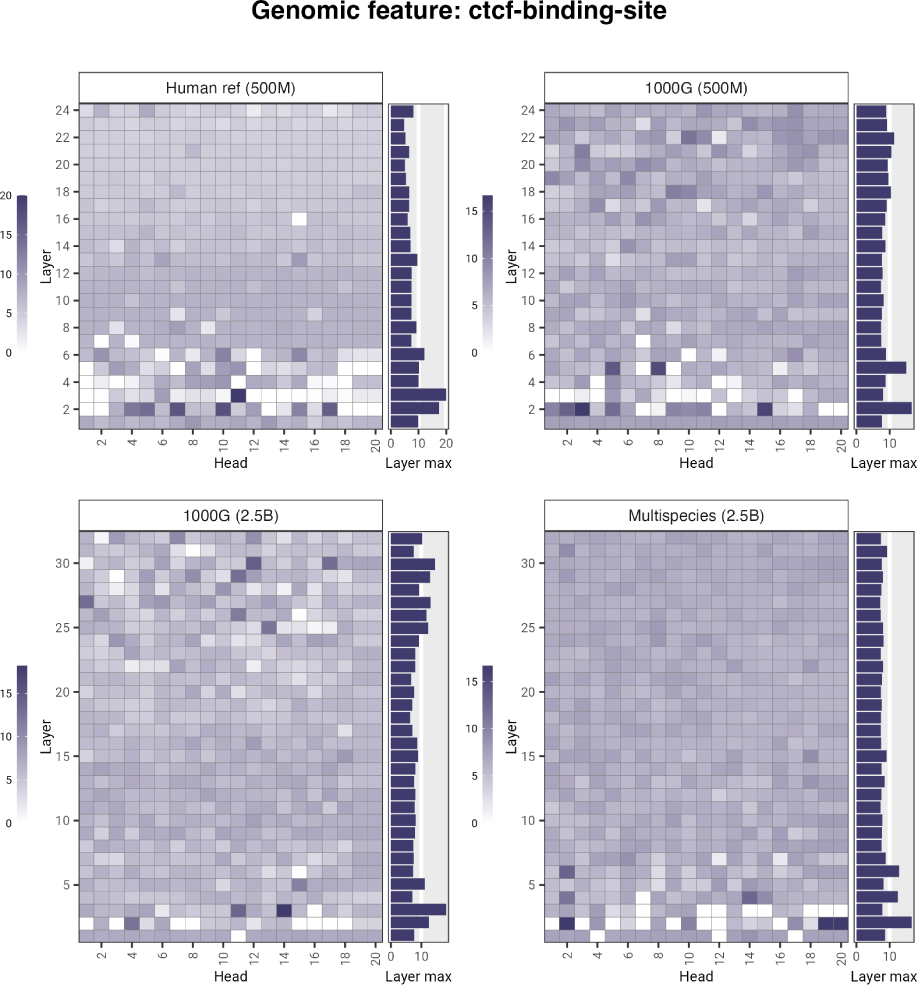
Nucleotide Transformer models’ attention percentages per head computed for CTCF binding sites.

**Supplementary Figure 12.**
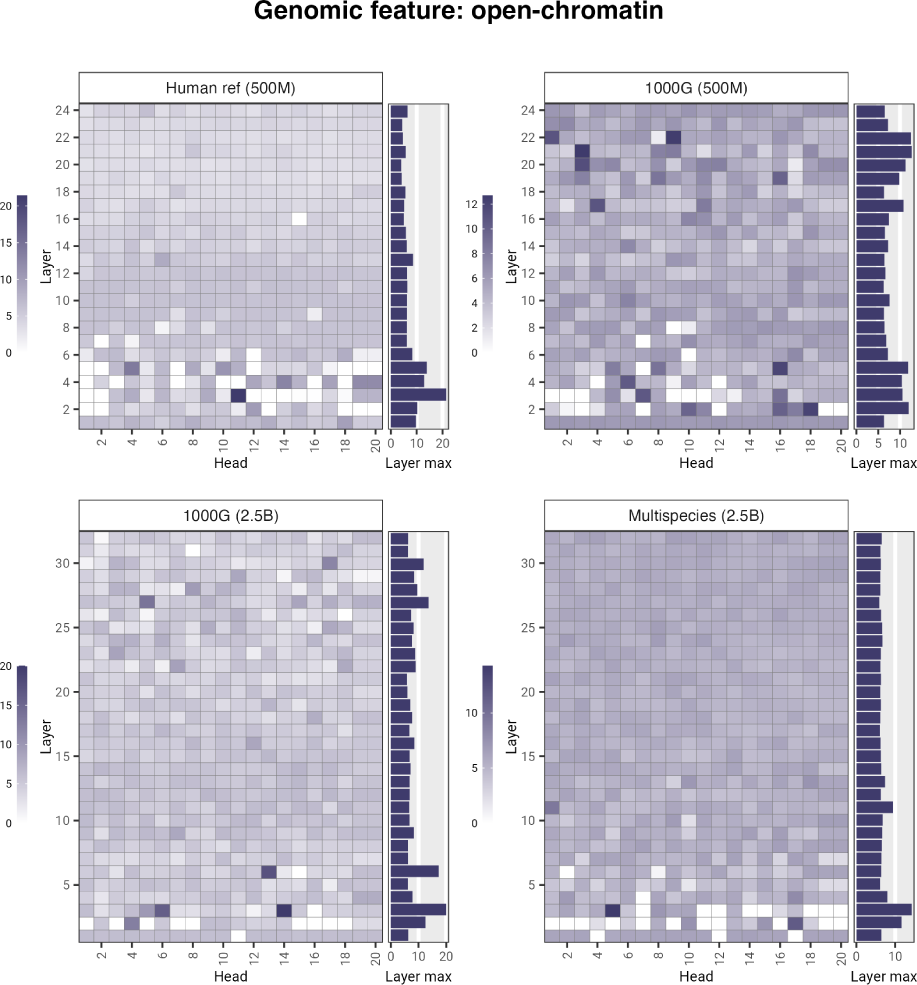
Nucleotide Transformer models’ attention percentages per head computed for open chromatin.

**Supplementary Figure 13.**
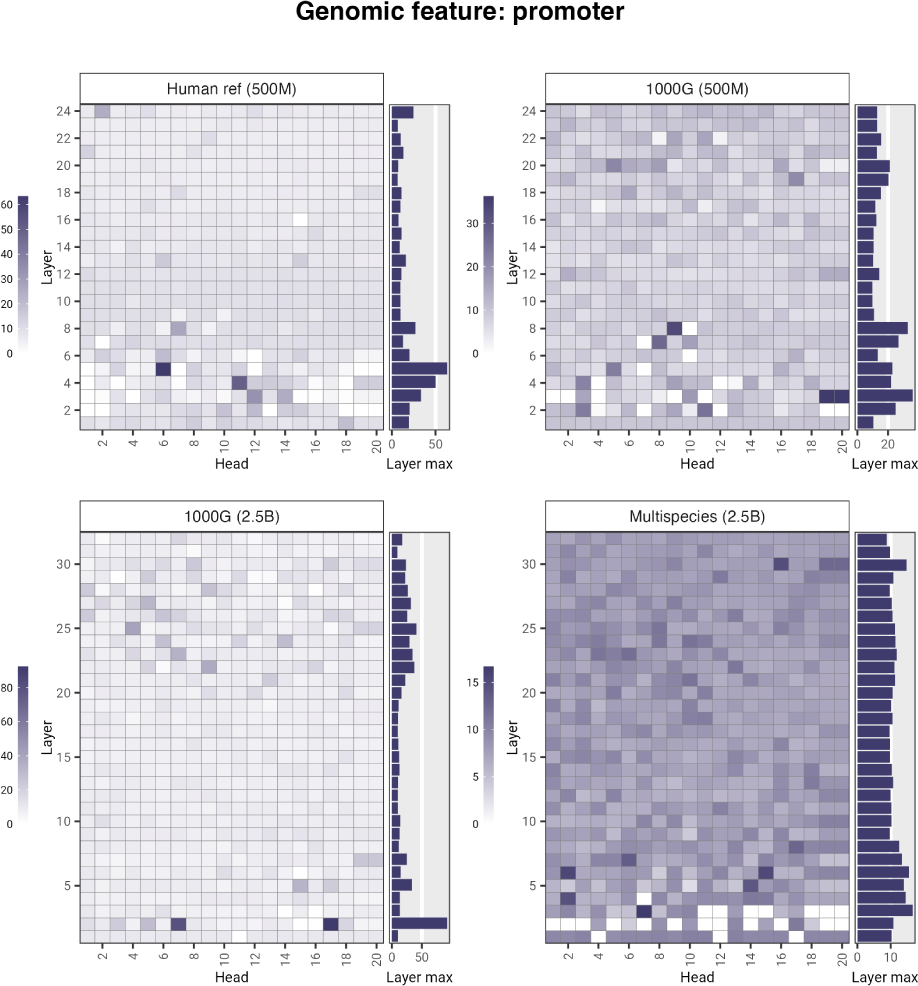
Nucleotide Transformer models’ attention percentages per head computed for promoters.

**Supplementary Figure 14.**
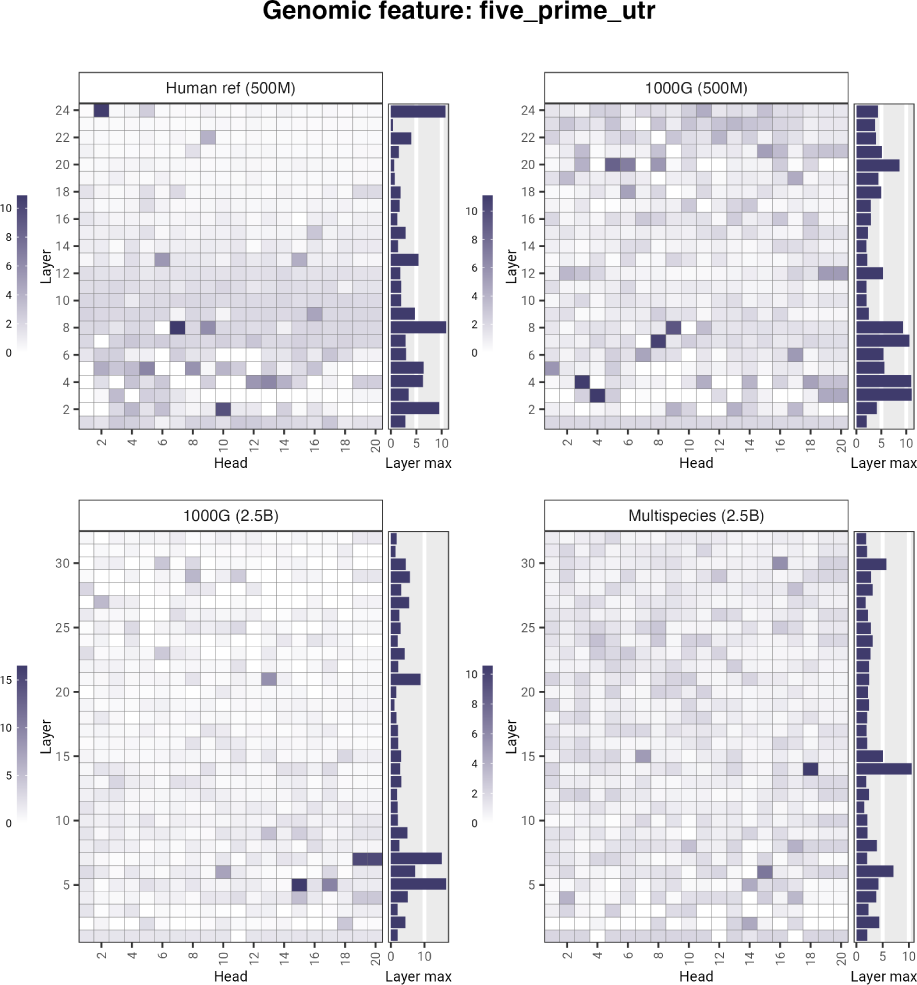
Nucleotide Transformer models’ attention percentages per head computed for 5’ UTR regions.

**Supplementary Figure 15.**
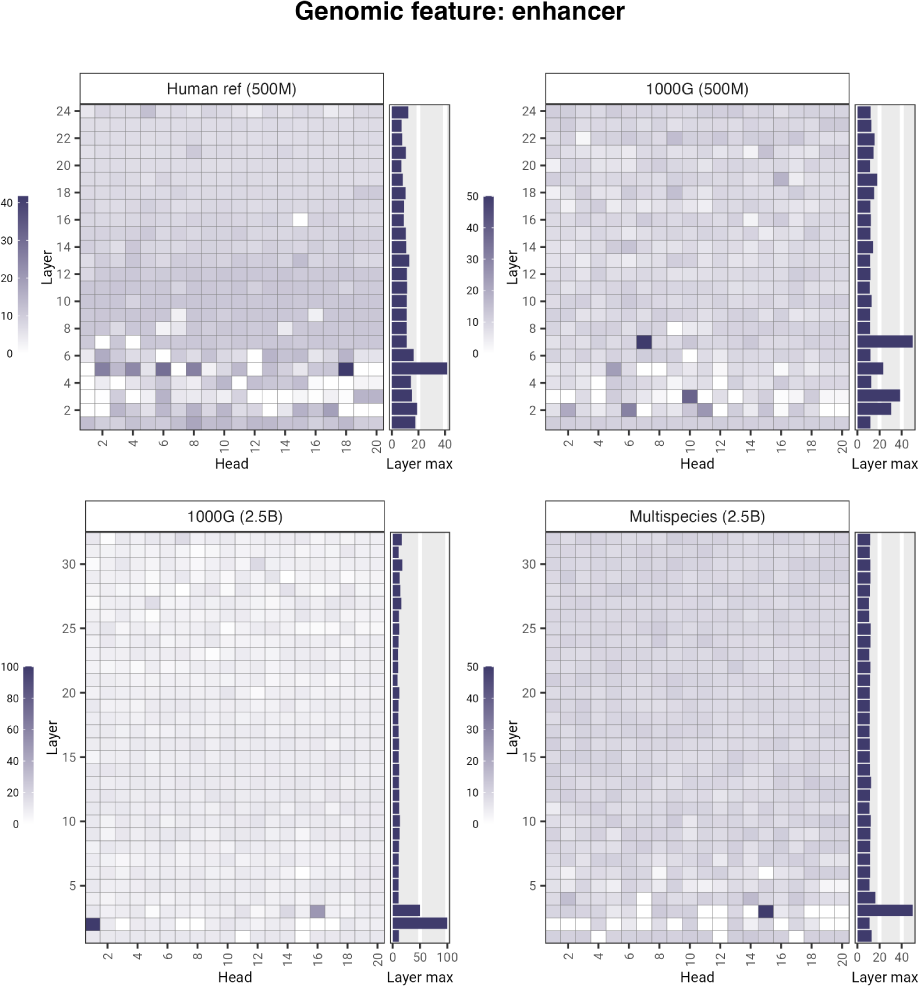
Nucleotide Transformer models’ attention percentages per head computed for enhancers.

**Supplementary Figure 16.**
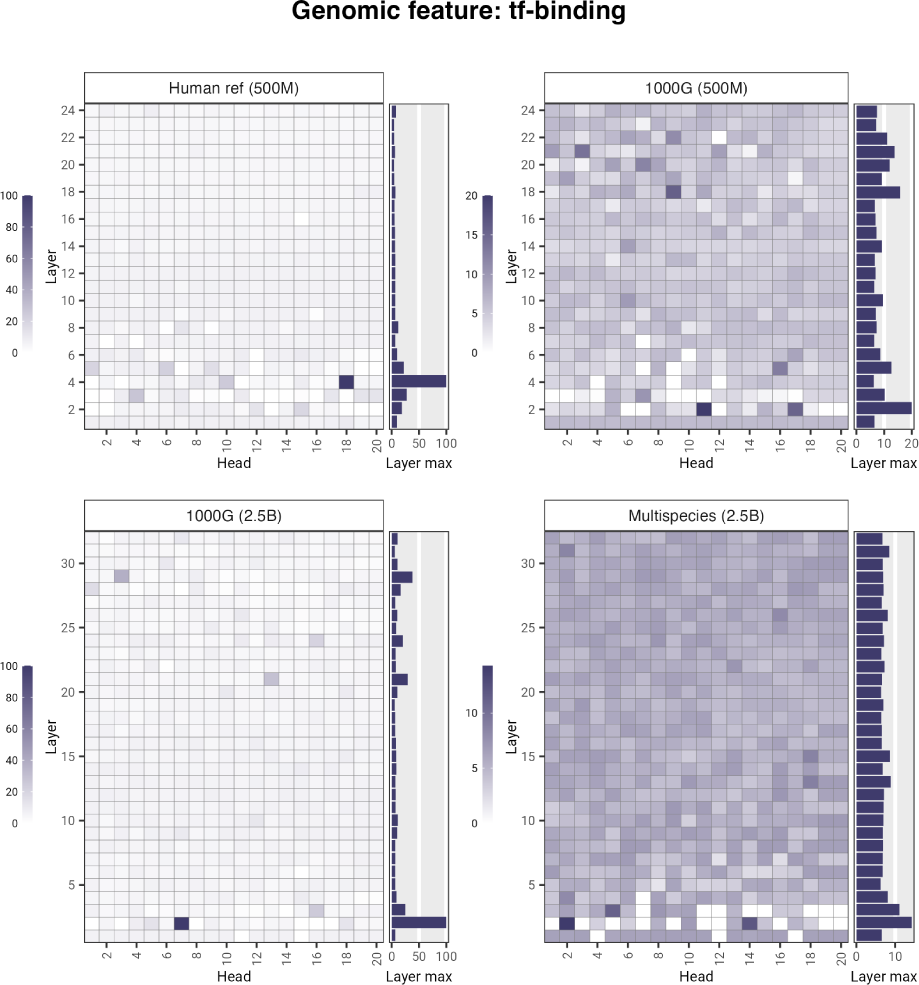
Nucleotide Transformer models’ attention percentages per head computed for transcription factor binding sites.

**Supplementary Figure 17.**
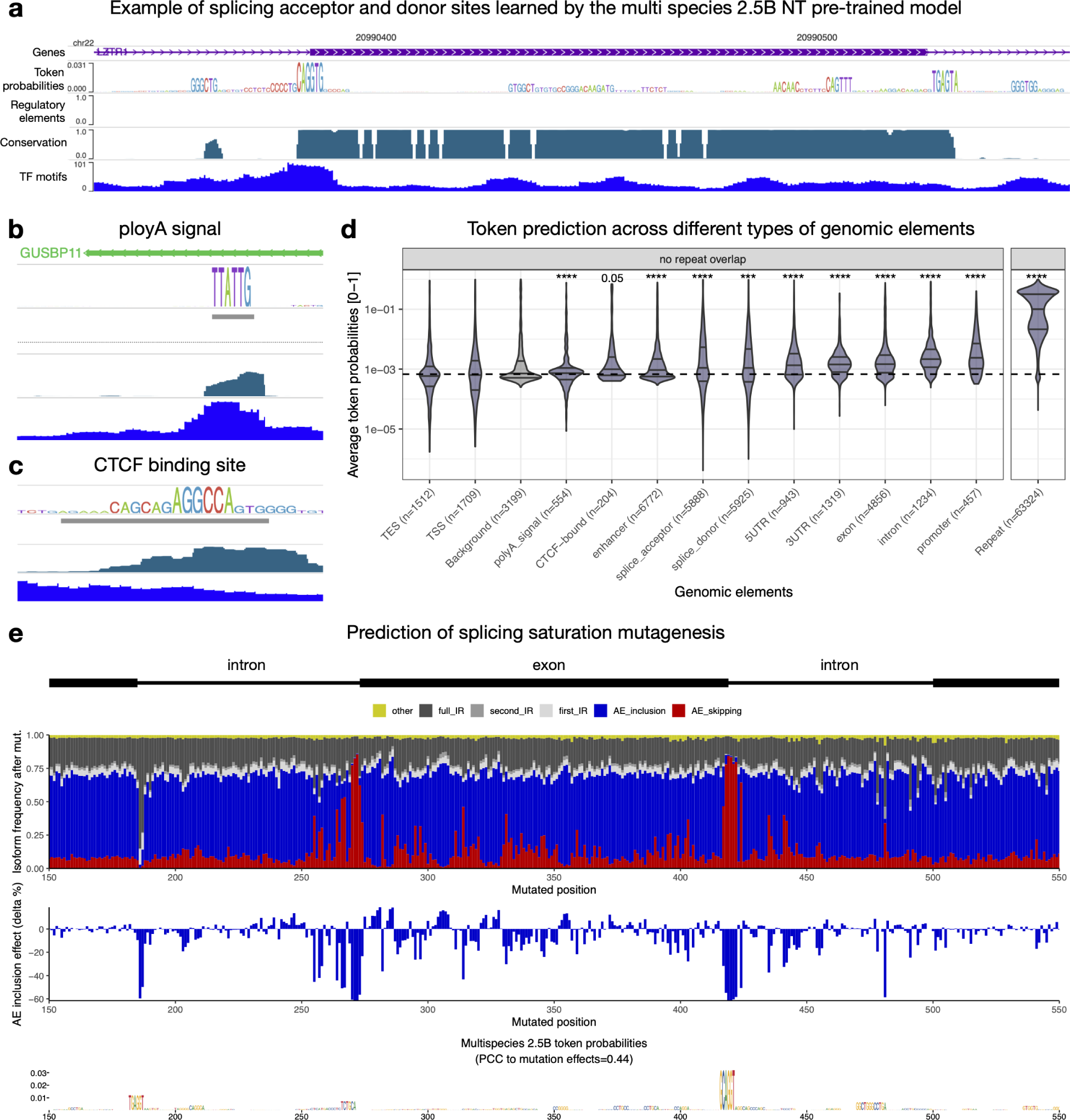
Interpretation of Nucleotide Transformer multispecies 2.5B pre-trained model at higher resolution. **a-c)** Genome browser screenshot depicting token probabilities for **(a)** an exon region of gene LZTR1 (chr22:20990296-20990630), **(b)** a polyA signal from GUSBP11 3’UTR (chr22:23638476-23638540), and **(c)** a CTCF binding site (chr22:22254952-22254981). ENCODE regulatory elements, conservation, and Jaspar TF motifs are also shown. **d)** Distribution of average token probability scores across different types of genomic elements that do not overlap repeat elements (left) or across repeat elements (right). Violins are shown with horizontal lines marking the median and lower and upper quartile. ****P < 0.0001, ***P < 0.001, **P < 0.01, *P < 0.05, (two-sided Wilcoxon rank-sum test). **e)** Top: experimentally measured impact of mutating every nucleotide position in the different splicing isoforms of exon 11 of gene MST1R. Middle: effect in Alternative Exon (AE) 11 inclusion as delta percentage. Bottom: token probability predictions by the multispecies 2.5B pre-trained model. Pearson Correlation Coefficient (PCC) between token predictions and AE11 inclusion mutation effects is shown.

**Supplementary Figure 18.**
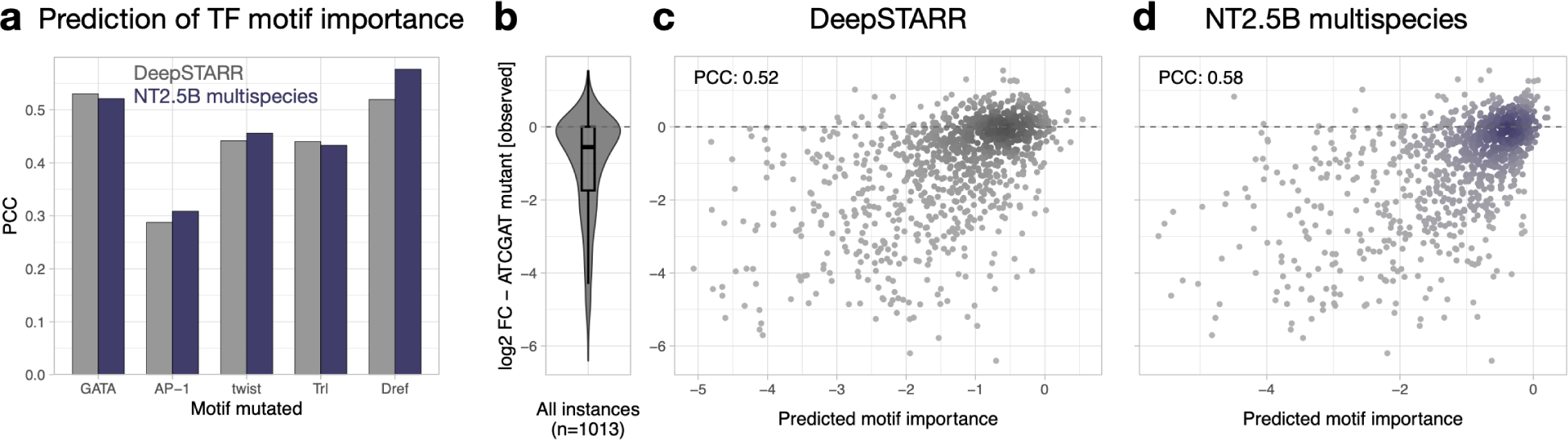
Prediction of transcription factor motif importance. **a)** Bar plots showing the Pearson Correlation Coefficient (PCC) between predicted (by DeepSTARR or the multispecies 2.5B full-finetuned model) and observed log2FC for mutating individual instances of each motif type. **b-d)** Distribution of experimentally measured enhancer activity log2FC after mutating 1,013 different Dref motif instances across enhancers (violin plot), compared with the log2FC predicted by **(c)** DeepSTARR or **(d)** the multispecies 2.5B model.

**Supplementary Figure 19.**
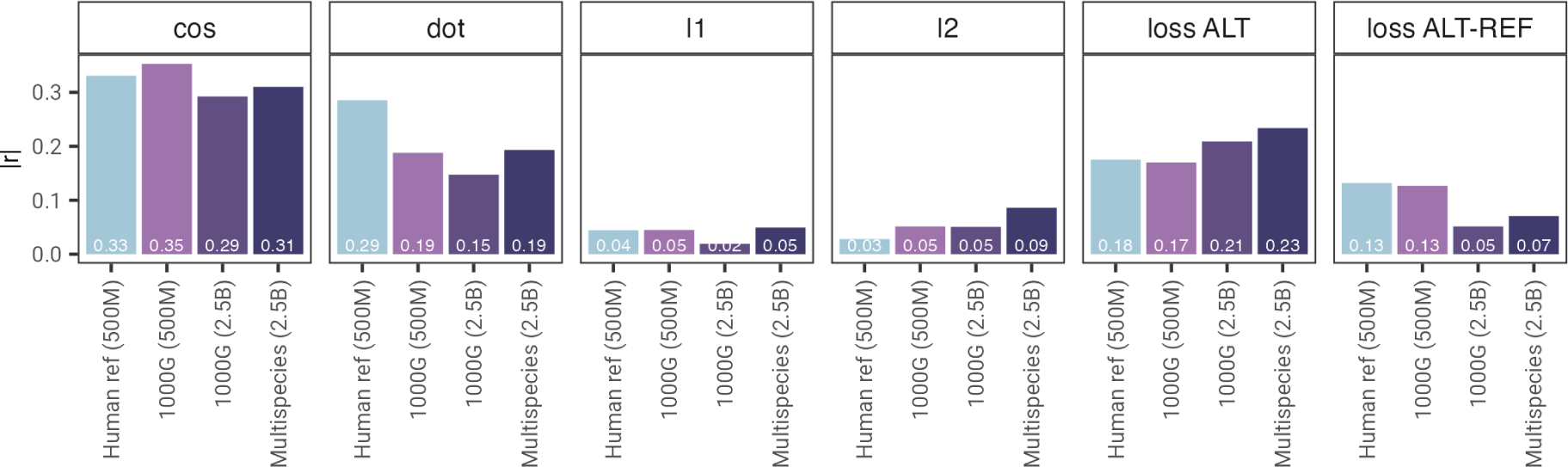
Performance of zero-shot based scores for SNPs annotations based on Ensembl Variant Effect Prediction (VEP). Performance is based on the Pearson correlation between the scores and severity as estimated by Ensembl.

## Supplementary Tables

**Supplementary Table 1.** Models hyperparameters.

|  | NT-1000G<br>(2.5B) | NT-Multispecies<br>(2.5B) | NT-1000G<br>(500M) | NT-HumanRef<br>(500M) | NT-Multispecies-v2<br>(500M) | NT-Multispecies-v2<br>(250M) | NT-Multispecies-v2<br>(100M) | NT-Multispecies-v2<br>(50M) |
| --- | --- | --- | --- | --- | --- | --- | --- | --- |
| alphabet | k-mers | k-mers | k-mers | k-mers | k-mers | k-mers | k-mers | k-mers |
| k_for_kmers | 6 | 6 | 6 | 6 | 6 | 6 | 6 | 6 |
| num_warmup_updates | 16000 | 16000 | 16000 | 16000 | 16000 | 16000 | 16000 | 16000 |
| warmup_init_lr | 5E-05 | 5E-05 | 5E-05 | 5E-05 | 5E-05 | 5E-05 | 5E-05 | 5E-05 |
| warmup_end_lr | 0.0001 | 0.0001 | 0.0001 | 0.0001 | 0.0001 | 0.0001 | 0.0001 | 0.0001 |
| training_set_proportion | 0.95 | 0.95 | 0.95 | 0.95 | 0.95 | 0.95 | 0.95 | 0.95 |
| masking_ratio | 0.15 | 0.15 | 0.15 | 0.15 | 0.15 | 0.15 | 0.15 | 0.15 |
| masking_prob | 0.8 | 0.8 | 0.8 | 0.8 | 0.8 | 0.8 | 0.8 | 0.8 |
| random_token_prob | 0 | 0.1 | 0 | 0.1 | 0.1 | 0.1 | 0.1 | 0.1 |
| alphabet_size | 4105 | 4105 | 4105 | 4105 | 4105 | 4105 | 4105 | 4105 |
| prepend_bos | True | True | True | True | True | True | True | True |
| append_eos | False | False | False | False | False | False | False | False |
| attention_heads | 20 | 20 | 20 | 20 | 16 | 16 | 16 | 16 |
| embed_dim | 2560 | 2560 | 1280 | 1280 | 1024 | 768 | 512 | 512 |
| ffn_embed_dim | 10240 | 10240 | 5120 | 5120 | 4096 | 3072 | 2048 | 2048 |
| num_layers | 32 | 32 | 24 | 24 | 29 | 24 | 22 | 12 |
| token_dropout | True | True | True | True | False | False | False | False |
| num_parameters | 2547800585 | 2547800585 | 485729545 | 485729545 | 496245771 | 233545995 | 96839179 | 54855179 |
| dataset | 1000G | Multispecies | 1000G | Human Reference | Multispecies | Multispecies | Multispecies | Multispecies |

**Supplementary Table 2.**
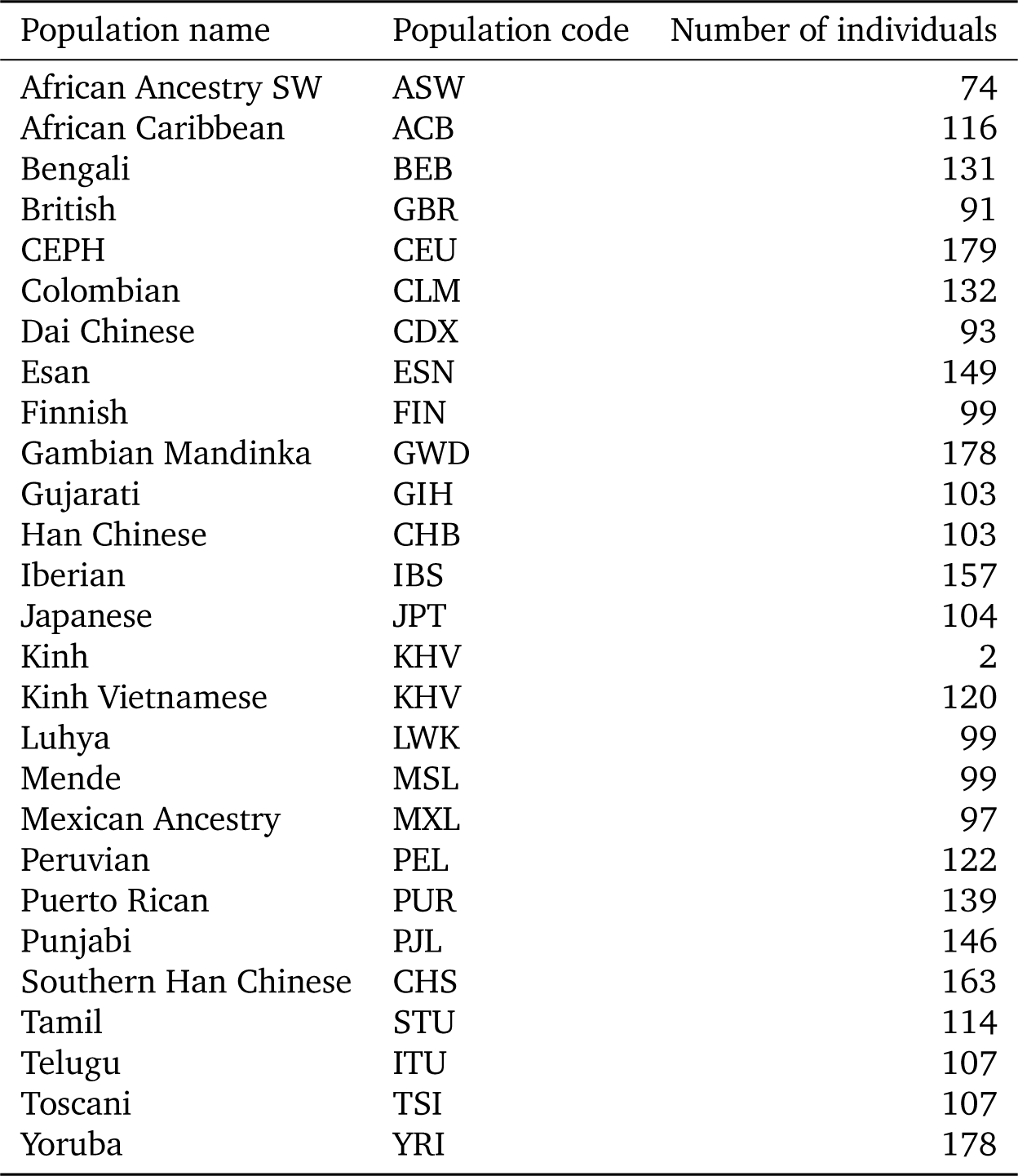
Number of individuals per population in the 1000G dataset.

**Supplementary Table 3.** Genomes in the multispecies dataset.

| Class | Number of species | Number of nucleotides (B) |
| --- | --- | --- |
| Bacteria | 667 | 17.1 |
| Fungi | 46 | 2.3 |
| Invertebrate | 39 | 20.8 |
| Protozoa | 10 | 0.5 |
| Mammalian Vertebrate | 31 | 69.8 |
| Other Vertebrate | 57 | 63.4 |

**Supplementary Table 4.** Model organisms genomes in the multispecies dataset.

| Class | Selected Species |
| --- | --- |
| Bacteria | Escherichia coli |
| Fungi | Saccharomyces cerevisiae |
| Invertebrate | Caenorhabditis elegans, Drosophila melanogaster |
| Protozoa | Plasmodium vivax, Plasmodium falciparum |
| Mammalian Vertebrate | Homo sapiens, Mus musculus, Rattus norvegicus |
| Other Vertebrate | Danio rerio, Xenopus tropicalis |

**Supplementary Table 5.** Nucleotide Transformer downstream tasks metadata.

|  | Num train sequences | Num test sequences | Max sequence length in bp | Reported Metric | Organisms |
| --- | --- | --- | --- | --- | --- |
| H2AFZ | 30000 | 3000 | 1000 | MCC | Human |
| H3K27ac | 30000 | 1616 | 1000 | MCC | Human |
| H3K27me3 | 30000 | 3000 | 1000 | MCC | Human |
| H3K36me3 | 30000 | 3000 | 1000 | MCC | Human |
| H3K4me1 | 30000 | 3000 | 1000 | MCC | Human |
| H3K4me2 | 30000 | 2138 | 1000 | MCC | Human |
| H3K4me3 | 17468 | 776 | 1000 | MCC | Human |
| H3K9ac | 23274 | 1004 | 1000 | MCC | Human |
| H3K9me3 | 27438 | 850 | 1000 | MCC | Human |
| H4K20me1 | 30000 | 2270 | 1000 | MCC | Human |
| Promoter all | 30000 | 1584 | 300 | MCC | Human |
| Promoter TATA | 30000 | 1372 | 300 | MCC | Human |
| Promoter non-TATA | 5062 | 212 | 300 | MCC | Human |
| Splice site all | 30000 | 3000 | 600 | MCC | Human |
| Splice donor | 30000 | 3000 | 600 | MCC | Human |
| Splice acceptor | 30000 | 3000 | 600 | MCC | Human |
| Enhancer | 30000 | 3000 | 400 | MCC | Human |
| Enhancer types | 30000 | 3000 | 400 | MCC | Human |

**Supplementary Table 6.** Downstream performance per task for all probing models. Every NT model was evaluated through probing and all “BPNet” models were fully trained CNN networks. Performance per task was calculated as the median of the 10 cross-validation folds (*±* standard deviation).

| Dataset Model | H2AFZ | H3K27ac | H3K27me3 | H3K36me3 | H3K4me1 | H3K4me2 |
| --- | --- | --- | --- | --- | --- | --- |
| Raw Probe | 0.302 (± 0.003) | 0.229 (± 0.006) | 0.379 (± 0.004) | 0.36 (± 0.003) | 0.287 (± 0.006) | 0.36 (± 0.005) |
| BPNet (original) | **0.473 (± 0.009)** | 0.296 (± 0.046) | 0.543 (± 0.009) | 0.548 (± 0.009) | 0.436 (± 0.008) | 0.427 (± 0.036) |
| BPNet (large) | **0.487 (± 0.014)** | 0.214 (± 0.037) | 0.551 (± 0.009) | **0.57 (± 0.009)** | **0.459 (± 0.012)** | 0.427 (± 0.025) |
| BPNet (X-large) | 0.475 (± 0.011) | 0.16 (± 0.042) | 0.542 (± 0.009) | 0.562 (± 0.011) | 0.457 (± 0.013) | 0.341 (± 0.038) |
| NT-HumanRef (500M) | 0.393 (± 0.01) | 0.422 (± 0.01) | 0.547 (± 0.007) | 0.516 (± 0.01) | 0.425 (± 0.014) | 0.453 (± 0.01) |
| NT-1000G (500M) | 0.404 (± 0.009) | 0.437 (± 0.007) | 0.535 (± 0.006) | 0.509 (± 0.005) | **0.438 (± 0.012)** | 0.515 (± 0.006) |
| NT-1000G (2.5B) | 0.425 (± 0.01) | **0.457 (± 0.011)** | **0.572 (± 0.009)** | **0.579 (± 0.009)** | **0.443 (± 0.015)** | **0.531 (± 0.007)** |
| NT-Multispecies (2.5B) | 0.422 (± 0.011) | **0.442 (± 0.019)** | 0.552 (± 0.009) | **0.569 (± 0.011)** | 0.427 (± 0.014) | 0.512 (± 0.007) |

| Dataset Model | H3K4me3 | H3K9ac | H3K9me3 | H4K20me1 | Enhancer | Enhancer types |
| --- | --- | --- | --- | --- | --- | --- |
| Raw Probe | 0.432 (± 0.008) | 0.322 (± 0.013) | 0.105 (± 0.01) | 0.446 (± 0.006) | 0.251 (± 0.003) | 0.233 (± 0.004) |
| BPNet (original) | 0.445 (± 0.047) | 0.336 (± 0.034) | 0.298 (± 0.03) | 0.531 (± 0.025) | **0.488 (± 0.009)** | **0.449 (± 0.006)** |
| BPNet (large) | 0.445 (± 0.049) | 0.298 (± 0.033) | 0.234 (± 0.037) | 0.525 (± 0.038) | **0.492 (± 0.008)** | **0.454 (± 0.008)** |
| BPNet (X-large) | 0.338 (± 0.076) | 0.235 (± 0.06) | 0.216 (± 0.085) | 0.411 (± 0.054) | **0.501 (± 0.013)** | **0.455 (± 0.012)** |
| NT-HumanRef (500M) | **0.594 (± 0.012)** | 0.458 (± 0.017) | 0.28 (± 0.021) | 0.608 (± 0.007) | 0.455 (± 0.011) | 0.423 (± 0.012) |
| NT-1000G (500M) | **0.592 (± 0.011)** | **0.523 (± 0.018)** | **0.381 (± 0.021)** | **0.622 (± 0.008)** | 0.47 (± 0.008) | 0.431 (± 0.005) |
| NT-1000G (2.5B) | **0.603 (± 0.018)** | **0.517 (± 0.014)** | **0.38 (± 0.031)** | **0.628 (± 0.012)** | 0.479 (± 0.007) | **0.439 (± 0.009)** |
| NT-Multispecies (2.5B) | **0.604 (± 0.016)** | **0.539 (± 0.015)** | **0.396 (± 0.011)** | **0.611 (± 0.011)** | 0.468 (± 0.009) | **0.446 (± 0.007)** |

**Supplementary Table 6** | Downstream performance per task for all probing models. Every NT model was evaluated through probing and all "BPNet" models were fully trained CNN networks. Performance per task was calculated as the median of the 10 cross-validation folds ( ± standard deviation).
| Dataset Model | Promoter all | Promoter non-TATA | Promoter TATA | Splice acceptor | Splice site all | Splice donor |
| --- | --- | --- | --- | --- | --- | --- |
| Raw Probe | 0.598 (± 0.002) | 0.619 (± 0.004) | 0.606 (± 0.007) | 0.136 (± 0.04) | 0.234 (± 0.003) | 0.395 (± 0.025) |
| BPNet (original) | 0.696 (± 0.026) | **0.717 (± 0.023)** | **0.848 (± 0.042)** | **0.859 (± 0.038)** | **0.793 (± 0.072)** | **0.92 (± 0.014)** |
| BPNet (large) | 0.672 (± 0.023) | 0.672 (± 0.043) | **0.826 (± 0.017)** | **0.925 (± 0.031)** | **0.865 (± 0.026)** | **0.93 (± 0.021)** |
| BPNet (X-large) | 0.579 (± 0.087) | 0.608 (± 0.061) | 0.607 (± 0.064) | 0.744 (± 0.01) | 0.745 (± 0.016) | 0.706 (± 0.009) |
| NT-HumanRef (500M) | 0.703 (± 0.007) | **0.725 (± 0.011)** | 0.662 (± 0.044) | 0.485 (± 0.012) | 0.414 (± 0.005) | 0.495 (± 0.01) |
| NT-1000G (500M) | 0.715 (± 0.011) | **0.732 (± 0.009)** | 0.696 (± 0.018) | 0.552 (± 0.006) | 0.355 (± 0.01) | 0.542 (± 0.01) |
| NT-1000G (2.5B) | **0.73 (± 0.009)** | **0.733 (± 0.013)** | 0.72 (± 0.024) | 0.594 (± 0.012) | 0.438 (± 0.008) | 0.621 (± 0.011) |
| NT-Multispecies (2.5B) | **0.743 (± 0.009)** | **0.747 (± 0.014)** | **0.774 (± 0.036)** | 0.664 (± 0.007) | 0.491 (± 0.01) | 0.687 (± 0.004) |

**Supplementary Table 7.** Downstream performance per task for all finetuned models. Every model was fine-tuned with PEFT, except for “BPNet” models that are fully trained CNN networks and “Raw Probe” that used probing strategy. Performance per task was calculated as the median of the 10 cross-validation folds (*±* standard deviation).

| Dataset Model | H2AFZ | H3K27ac | H3K27me3 | H3K36me3 | H3K4me1 | H3K4me2 |
| --- | --- | --- | --- | --- | --- | --- |
| Raw Probe | 0.302 (± 0.003) | 0.229 (± 0.006) | 0.379 (± 0.004) | 0.36 (± 0.003) | 0.287 (± 0.006) | 0.36 (± 0.005) |
| BPNet (original) | 0.473 (± 0.009) | 0.296 (± 0.046) | 0.543 (± 0.009) | 0.548 (± 0.009) | 0.436 (± 0.008) | 0.427 (± 0.036) |
| BPNet (large) | 0.487 (± 0.014) | 0.214 (± 0.037) | 0.551 (± 0.009) | 0.57 (± 0.009) | 0.459 (± 0.012) | 0.427 (± 0.025) |
| BPNet (X-large) | 0.475 (± 0.011) | 0.16 (± 0.042) | 0.542 (± 0.009) | 0.562 (± 0.011) | 0.457 (± 0.013) | 0.341 (± 0.038) |
| DNABERT-2 | 0.49 (± 0.013) | 0.491 (± 0.01) | **0.599 (± 0.01)** | **0.637 (± 0.007)** | **0.49 (± 0.008)** | 0.558 (± 0.013) |
| Enformer | **0.522 (± 0.019)** | **0.52 (± 0.015)** | 0.552 (± 0.007) | 0.567 (± 0.017) | **0.504 (± 0.021)** | **0.626 (± 0.015)** |
| HyenaDNA-1KB | 0.455 (± 0.015) | 0.423 (± 0.017) | 0.541 (± 0.018) | 0.543 (± 0.01) | 0.43 (± 0.014) | 0.521 (± 0.024) |
| HyenaDNA-32KB | 0.467 (± 0.012) | 0.421 (± 0.01) | 0.55 (± 0.009) | 0.553 (± 0.011) | 0.423 (± 0.016) | 0.515 (± 0.018) |
| NT 500M Randomly Initialized | 0.281 (± 0.017) | 0.233 (± 0.015) | 0.364 (± 0.021) | 0.305 (± 0.004) | 0.228 (± 0.018) | 0.317 (± 0.021) |
| NT 2.5B Randomly Initialized | 0.291 (± 0.011) | 0.257 (± 0.02) | 0.379 (± 0.009) | 0.324 (± 0.014) | 0.271 (± 0.02) | 0.33 (± 0.014) |
| NT-HumanRef (500M) | 0.465 (± 0.011) | 0.457 (± 0.01) | 0.589 (± 0.009) | 0.594 (± 0.004) | 0.468 (± 0.007) | 0.527 (± 0.011) |
| NT-1000G (500M) | 0.464 (± 0.012) | 0.458 (± 0.012) | 0.591 (± 0.007) | 0.581 (± 0.009) | 0.466 (± 0.006) | 0.528 (± 0.011) |
| NT-1000G (2.5B) | 0.478 (± 0.012) | **0.486 (± 0.023)** | **0.603 (± 0.009)** | **0.632 (± 0.008)** | **0.491 (± 0.015)** | 0.569 (± 0.014) |
| NT-Multispecies (2.5B) | **0.503 (± 0.01)** | 0.481 (± 0.02) | **0.593 (± 0.016)** | **0.635 (± 0.016)** | **0.481 (± 0.012)** | 0.552 (± 0.022) |
| NT-Multispecies-v2 (50M) | **0.501 (± 0.012)** | 0.483 (± 0.012) | 0.595 (± 0.004) | 0.607 (± 0.006) | **0.478 (± 0.012)** | 0.549 (± 0.014) |
| NT-Multispecies-v2 (100M) | **0.492 (± 0.012)** | 0.487 (± 0.016) | **0.595 (± 0.013)** | 0.617 (± 0.006) | **0.485 (± 0.011)** | 0.551 (± 0.01) |
| NT-Multispecies-v2 (250M) | **0.513 (± 0.017)** | **0.497 (± 0.014)** | **0.6 (± 0.009)** | **0.636 (± 0.012)** | **0.491 (± 0.006)** | 0.57 (± 0.009) |
| NT-Multispecies-v2 (500M) | **0.493 (± 0.01)** | 0.492 (± 0.012) | **0.613 (± 0.01)** | **0.642 (± 0.015)** | **0.497 (± 0.009)** | 0.564 (± 0.012) |

| Dataset Model | H3K4me3 | H3K9ac | H3K9me3 | H4K20me1 | Enhancer | Enhancer types |
| --- | --- | --- | --- | --- | --- | --- |
| Raw Probe | 0.432 (± 0.008) | 0.322 (± 0.013) | 0.105 (± 0.01) | 0.446 (± 0.006) | 0.251 (± 0.003) | 0.233 (± 0.004) |
| BPNet (original) | 0.445 (± 0.047) | 0.336 (± 0.034) | 0.298 (± 0.03) | 0.531 (± 0.025) | 0.488 (± 0.009) | 0.449 (± 0.006) |
| BPNet (large) | 0.445 (± 0.049) | 0.298 (± 0.033) | 0.234 (± 0.037) | 0.525 (± 0.038) | 0.492 (± 0.008) | 0.454 (± 0.008) |
| BPNet (X-large) | 0.338 (± 0.076) | 0.235 (± 0.06) | 0.216 (± 0.085) | 0.411 (± 0.054) | 0.501 (± 0.013) | 0.455 (± 0.012) |
| DNABERT-2 | **0.646 (± 0.008)** | **0.564 (± 0.013)** | 0.443 (± 0.025) | **0.655 (± 0.011)** | 0.517 (± 0.011) | 0.476 (± 0.009) |
| Enformer | **0.635 (± 0.019)** | **0.593 (± 0.02)** | 0.453 (± 0.016) | 0.606 (± 0.016) | **0.614 (± 0.01)** | **0.573 (± 0.013)** |
| HyenaDNA-1KB | 0.596 (± 0.015) | 0.484 (± 0.022) | 0.375 (± 0.026) | 0.58 (± 0.009) | 0.475 (± 0.006) | 0.441 (± 0.01) |
| HyenaDNA-32KB | 0.603 (± 0.02) | 0.487 (± 0.025) | 0.419 (± 0.03) | 0.59 (± 0.007) | 0.476 (± 0.021) | 0.445 (± 0.009) |
| NT 500M Randomly Initialized | 0.452 (± 0.031) | 0.329 (± 0.035) | 0.088 (± 0.015) | 0.431 (± 0.012) | 0.251 (± 0.012) | 0.215 (± 0.022) |
| NT 2.5B Randomly Initialized | 0.463 (± 0.023) | 0.299 (± 0.021) | 0.097 (± 0.035) | 0.436 (± 0.01) | 0.276 (± 0.012) | 0.225 (± 0.021) |
| NT-HumanRef (500M) | 0.622 (± 0.013) | 0.524 (± 0.013) | 0.433 (± 0.009) | 0.634 (± 0.013) | 0.515 (± 0.019) | 0.477 (± 0.014) |
| NT-1000G (500M) | 0.609 (± 0.011) | 0.515 (± 0.018) | 0.415 (± 0.019) | 0.634 (± 0.01) | 0.505 (± 0.009) | 0.459 (± 0.011) |
| NT-1000G (2.5B) | 0.615 (± 0.017) | 0.529 (± 0.012) | **0.483 (± 0.013)** | **0.659 (± 0.008)** | 0.504 (± 0.009) | 0.469 (± 0.005) |
| NT-Multispecies (2.5B) | 0.618 (± 0.015) | 0.527 (± 0.017) | 0.447 (± 0.018) | **0.65 (± 0.014)** | 0.527 (± 0.012) | 0.484 (± 0.012) |
| NT-Multispecies-v2 (50M) | **0.627 (± 0.013)** | 0.538 (± 0.011) | 0.43 (± 0.018) | 0.637 (± 0.008) | 0.511 (± 0.006) | 0.459 (± 0.006) |
| NT-Multispecies-v2 (100M) | **0.633 (± 0.015)** | 0.538 (± 0.015) | 0.445 (± 0.017) | **0.648 (± 0.008)** | 0.507 (± 0.009) | 0.465 (± 0.009) |
| NT-Multispecies-v2 (250M) | **0.64 (± 0.009)** | **0.565 (± 0.021)** | **0.467 (± 0.016)** | **0.652 (± 0.006)** | 0.525 (± 0.01) | 0.492 (± 0.01) |
| NT-Multispecies-v2 (500M) | **0.636 (± 0.016)** | 0.539 (± 0.013) | **0.475 (± 0.014)** | **0.658 (± 0.008)** | 0.524 (± 0.013) | 0.498 (± 0.009) |

**Supplementary Table 7** | Downstream performance per task for all finetuned models.
| Dataset Model | Promoter all | Promoter non-TATA | Promoter TATA | Splice acceptor | Splice site all | Splice donor |
| --- | --- | --- | --- | --- | --- | --- |
| Raw Probe | 0.598 (± 0.002) | 0.619 (± 0.004) | 0.606 (± 0.007) | 0.136 (± 0.04) | 0.234 (± 0.003) | 0.395 (± 0.025) |
| BPNet (original) | 0.696 (± 0.026) | 0.717 (± 0.023) | 0.848 (± 0.042) | 0.859 (± 0.038) | 0.793 (± 0.072) | 0.92 (± 0.014) |
| BPNet (large) | 0.672 (± 0.023) | 0.672 (± 0.043) | 0.826 (± 0.017) | **0.925 (± 0.031)** | 0.865 (± 0.026) | 0.93 (± 0.021) |
| BPNet (X-large) | 0.579 (± 0.087) | 0.608 (± 0.061) | 0.607 (± 0.064) | 0.744 (± 0.01) | 0.745 (± 0.016) | 0.706 (± 0.009) |
| DNABERT-2 | 0.754 (± 0.009) | **0.769 (± 0.009)** | 0.784 (± 0.036) | 0.837 (± 0.006) | 0.855 (± 0.005) | 0.861 (± 0.004) |
| Enformer | 0.745 (± 0.012) | 0.763 (± 0.012) | 0.793 (± 0.026) | 0.749 (± 0.007) | 0.739 (± 0.011) | 0.78 (± 0.007) |
| HyenaDNA-1KB | 0.693 (± 0.016) | 0.723 (± 0.013) | 0.648 (± 0.044) | 0.815 (± 0.049) | 0.854 (± 0.053) | 0.943 (± 0.024) |
| HyenaDNA-32KB | 0.698 (± 0.011) | 0.729 (± 0.009) | 0.666 (± 0.041) | 0.808 (± 0.009) | 0.907 (± 0.018) | 0.915 (± 0.047) |
| NT 500M Randomly Initialized | 0.575 (± 0.01) | 0.581 (± 0.025) | 0.39 (± 0.044) | 0.207 (± 0.031) | 0.363 (± 0.038) | 0.596 (± 0.057) |
| NT 2.5B Randomly Initialized | 0.589 (± 0.014) | 0.597 (± 0.022) | 0.465 (± 0.04) | 0.258 (± 0.049) | 0.421 (± 0.024) | 0.641 (± 0.045) |
| NT-HumanRef (500M) | 0.734 (± 0.013) | 0.738 (± 0.008) | 0.831 (± 0.022) | 0.941 (± 0.004) | 0.939 (± 0.003) | 0.952 (± 0.003) |
| NT-1000G (500M) | 0.727 (± 0.004) | 0.743 (± 0.012) | 0.855 (± 0.041) | 0.933 (± 0.007) | 0.939 (± 0.004) | 0.952 (± 0.004) |
| NT-1000G (2.5B) | 0.708 (± 0.008) | 0.758 (± 0.007) | 0.802 (± 0.03) | **0.952 (± 0.004)** | 0.956 (± 0.004) | 0.963 (± 0.001) |
| NT-Multispecies (2.5B) | **0.761 (± 0.009)** | **0.773 (± 0.01)** | **0.944 (± 0.016)** | **0.958 (± 0.003)** | **0.964 (± 0.003)** | **0.97 (± 0.002)** |
| NT-Multispecies-v2 (50M) | 0.731 (± 0.01) | 0.756 (± 0.01) | 0.747 (± 0.043) | 0.922 (± 0.006) | 0.928 (± 0.005) | 0.915 (± 0.006) |
| NT-Multispecies-v2 (100M) | 0.753 (± 0.005) | **0.766 (± 0.014)** | 0.826 (± 0.019) | 0.947 (± 0.003) | **0.96 (± 0.005)** | 0.947 (± 0.008) |
| NT-Multispecies-v2 (250M) | **0.774 (± 0.013)** | **0.785 (± 0.01)** | 0.87 (± 0.019) | **0.95 (± 0.008)** | **0.965 (± 0.003)** | **0.967 (± 0.004)** |
| NT-Multispecies-v2 (500M) | **0.778 (± 0.012)** | **0.774 (± 0.015)** | 0.878 (± 0.021) | 0.944 (± 0.005) | **0.967 (± 0.004)** | **0.96 (± 0.01)** |

**Supplementary Table 8.** Hyperparameters used to train the probing model with the best cross-validation performance for each of the downstream task, along with the model and the layer used to compute the corresponding embeddings. The best probe method for the tasks Enhancer types and Splice donor was the logistic regression with default hyperparameters.

|  | Model | Layer probed | Number of hidden layers | Hidden layer dimension | Batch size | Learning rate |
| --- | --- | --- | --- | --- | --- | --- |
| Enhancer | 2.5B 1000G | 17 | 1 | 256 | 500 | 1e-4 |
| H4K20me1 | 2.5B 1000G | 17 | 1 | 256 | 500 | 2e-4 |
| H2AFZ | 2.5B 1000G | 17 | 1 | 256 | 500 | 1e-4 |
| H3K4me1 | 2.5B 1000G | 14 | 2 | 512 | 500 | 1e-4 |
| H3K9ac | 2.5B MS | 10 | 1 | 256 | 500 | 1e-4 |
| Promoter all | 2.5B MS | 10 | 1 | 512 | 500 | 3e-4 |
| Promoter non-TATA | 2.5B MS | 10 | 1 | 256 | 500 | 1e-4 |
| H3K36me3 | 2.5B 1000G | 17 | 1 | 256 | 500 | 1e-4 |
| Enhancer types | 2.5B MS | 6 | 1 | 256 | 500 | 1e-4 |
| H3K27me3 | 2.5B 1000G | 17 | 1 | 256 | 500 | 1e-4 |
| Splice acceptor | 2.5B MS | 10 | 1 | 256 | 500 | 1e-4 |
| H3K4me3 | 2.5B MS | 17 | 1 | 256 | 500 | 1e-4 |
| Promoter TATA | 2.5B MS | 10 | 2 | 512 | 500 | 5e-4 |
| Splice donor | 2.5B MS | 10 | 1 | 512 | 500 | 1e-4 |
| H3K4me2 | 2.5B 1000G | 17 | 1 | 256 | 500 | 1e-4 |
| H3K9me3 | 2.5B MS | 10 | 1 | 256 | 500 | 1e-4 |
| H3K27ac | 2.5B 1000G | 17 | 1 | 256 | 500 | 1e-4 |
| Splice site all | 2.5B MS | 14 | 1 | 512 | 652 | 1e-4 |

**Supplementary Table 9.** Normalized summed downstream performance of all models, sorted by performance. These values are obtained by averaging the median MCC score obtained by each model on the three categories of downstream tasks: Chromatin Profiles, Regulatory elements and Splicing. Information about model type and probing/fine-tuning approach is also provided.

| Model name | Model type | Approach | Pre-training | Average performance |
| --- | --- | --- | --- | --- |
| NT 500M Randomly Initialized | Control | Training from scratch | NA | 0.355 |
| Raw Probe | Raw Probe | Probing | Self-supervised | 0.391 |
| NT 2.5B Randomly Initialized | Control | Training from scratch | NA | 0.399 |
| DNABERT-1 | DNABERT-1 | Finetuning | Self-supervised | 0.511 |
| NT-HumanRef (500M) | NT-v1 | Probing | Self-supervised | 0.534 |
| BPNet (X-large) | BPNet | control | NA | 0.566 |
| NT-1000G (500M) | NT-v1 | Probing | Self-supervised | 0.583 |
| NT-1000G (2.5B) | NT-v1 | Probing | Self-supervised | 0.613 |
| NT-Multispecies (2.5B) | NT-v1 | Probing | Self-supervised | 0.644 |
| BPNet (original) | BPNet | control | NA | 0.665 |
| HyenaDNA-1KB | HyenaDNA | Finetuning | Self-supervised | 0.668 |
| BPNet (large) | BPNet | control | NA | 0.683 |
| Enformer | Enformer | Finetuning | Supervised | 0.684 |
| HyenaDNA-32KB | HyenaDNA | Finetuning | Self-supervised | 0.691 |
| DNABERT-2 | DNABERT-2 | Finetuning | Self-supervised | 0.723 |
| NT-1000G (500M) | NT-v1 | Finetuning | Self-supervised | 0.729 |
| NT-Multispecies-v2 (50M) | NT-v2 | Finetuning | Self-supervised | 0.732 |
| NT-HumanRef (500M) | NT-v1 | Finetuning | Self-supervised | 0.733 |
| NT-1000G (2.5B) | NT-v1 | Finetuning | Self-supervised | 0.738 |
| NT-Multispecies-v2 (100M) | NT-v2 | Finetuning | Self-supervised | 0.748 |
| NT-Multispecies (2.5B) | NT-v1 | Finetuning | Self-supervised | 0.755 |
| NT-Multispecies-v2 (500M) | NT-v2 | Finetuning | Self-supervised | 0.762 |
| NT-Multispecies-v2 (250M) | NT-v2 | Finetuning | Self-supervised | **0.769** |

**Supplementary Table 10.** Proportions of the genomic elements in the dataset used for attention maps and embedding space visualization experiments.

|  | Percentage of sequences<br>containing this element | Mean Percentage of Sequence Length<br>belonging to this element |
| --- | --- | --- |
| Enhancer | 10.18% | 11.09% |
| Open chromatin | 11.32% | 6.16% |
| 3' UTR | 11.37% | 4.30% |
| TF binding | 11.28% | 6.38% |
| Intron | 10.45% | 11.08% |
| Exon | 11.87% | 3.18% |
| 5'UTR | 11.84% | 1.99% |
| Promoter | 10.95% | 9.78% |
| CTCF Binding Site | 10.69% | 7.36% |

**Supplementary Table 11.**
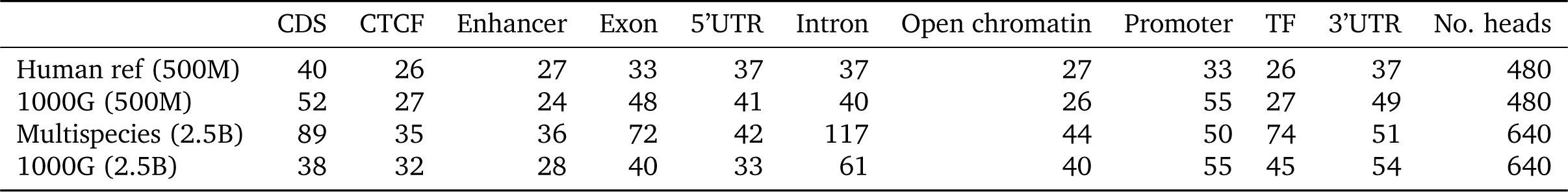
Number of significant attention heads capturing gene features and regulatory elements. Note that the number of total heads varies between the 500M and 2.5B models.

**Supplementary Table 12.**
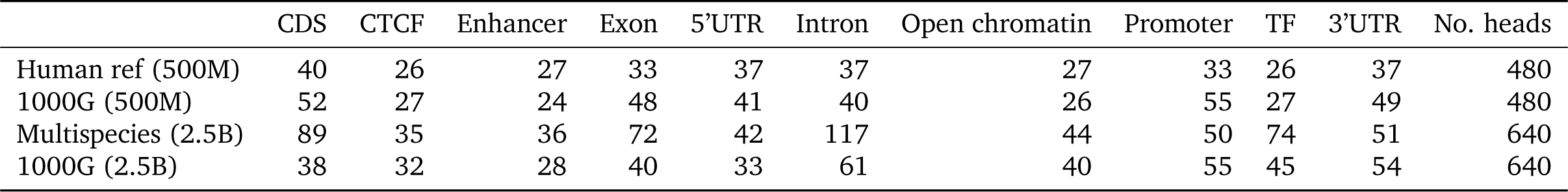
Number of significant attention heads capturing gene features and regulatory elements. Note that the number of total heads varies between the 500M and 2.5B models.

## Footnotes

1 https://huggingface.co/spaces/InstaDeepAI/nucleotide_transformer_benchmar^k^

2 https://huggingface.co/spaces/InstaDeepAI/nucleotide_transformer_benchmar^k^

3 https://www.ncbi.nlm.nih.gov/assembly/GCF_000001405.2^6^

4 http://ftp.1000genomes.ebi.ac.uk/vol1/ftp/data_collections/1000G_2504_high_coverage/working/20201028_3202_phased/

5 https://www.ncbi.nlm.nih.gov^/^

6 https://jax.readthedocs.io/en/latest/_autosummary/jax.pmap.htm^l^

7 https://developer.nvidia.com/ncc^l^

8 http://deepsea.princeton.edu/media/code/deepsea_train_bundle.v0.9.tar.g^z^

9 https://basespace.illumina.com/projects/66029966^/^

10 https://zenodo.org/record/5502060#.Y9qO7hzMLU^c^

11 https://shorturl.at/jov2^8^

12 https://www.ensembl.org/info/data/ftp/index.htm^l^

13 https://www.gencodegenes.org/human^/^

14 https://api.wenglab.org/screen_v13/fdownloads/GRCh38-ccREs.be^d^

